# Deletion of Mcpip1 in Mcpip1^AlbKO^ mice recapitulates the phenotype of human primary biliary cholangitis

**DOI:** 10.1101/2020.09.05.250522

**Authors:** Jerzy Kotlinowski, Tomasz Hutsch, Izabela Czyzynska-Cichon, Marta Wadowska, Natalia Pydyn, Agnieszka Jasztal, Agnieszka Kij, Ewelina Dobosz, Maciej Lech, Katarzyna Miekus, Ewelina Pośpiech, Mingui Fu, Jolanta Jura, Joanna Koziel, Stefan Chlopicki

## Abstract

**Background & Aims:** Primary biliary cholangitis (PBC) is an autoimmune disease characterized by progressive destruction of the intrahepatic bile ducts. The immunopathology of PBC involves excessive inflammation; therefore, negative regulators of inflammatory response, such as Monocyte Chemoattractant Protein-1-Induced Protein-1 (MCPIP1, *alias* Regnase1) may play important roles in the development of PBC. The aim of this work was to verify whether Mcpip1 expression protects against development of PBC.

**Methods:** Genetic deletion of *Zc3h12a* was used to characterize the role of Mcpip1 in the pathogenesis of PBC. 6-52-week-old Mcpip1^fl/fl^ and Mcpip1^AlbKO^ mice were used for immunohistochemical, biochemical and molecular tests.

**Results:** We found that Mcpip1 deficiency in the liver recapitulates most of the features of human PBC, in contrast to mice with Mcpip1 deficiency in myeloid cells (Mcpip1^LysMKO^ mice), which present with robust myeloid cell-driven systemic inflammation. In Mcpip1^AlbKO^ livers, intrahepatic bile ducts displayed proliferative changes with inflammatory infiltration, bile duct destruction, and fibrosis leading to cholestasis. In plasma, increased concentrations of IgG, IgM, and AMA autoantibodies (anti-PDC-E2) were detected. Interestingly, the phenotype of Mcpip1^AlbKO^ mice was robust in 6-week-old and 52-week-old mice, but milder in 12-24-week-old mice, suggesting early prenatal origin of the phenotype and age-dependent progression of the disease. Hepatic transcriptome analysis of 6-week-old and 24-week-old Mcpip1^AlbKO^ mice showed 812 and 8 differentially expressed genes (DEGs), respectively, compared with age-matched control mice, and revealed a distinct set of genes compared to those previously associated with development of PBC.

**Conclusions:** The phenotype of Mcpip1^AlbKO^ mice recapitulates most of the features of human PBC, and demonstrates early prenatal origin and age-dependent progression of PBC. Therefore, Mcpip1^AlbKO^ mice provide a unique model for the study of PBC.

**Lay summary:** Deletion of hepatic Mcpip1 in Mcpip1^AlbKO^ mice leads to development of PBC that recapitulates phenotype of human patients. These animals, show early prenatal origin and age-dependent progression of the disease. Thus, Mcpip1^AlbKO^ mice provide a unique model for studying PBC.

## Introduction

Primary biliary cholangitis (PBC) is an autoimmune, chronic cholestatic liver disease, characterized by a slow, progressive destruction of the intrahepatic bile ducts. Features of the disease include inflammation leading to the destruction of small- and medium-sized intrahepatic bile ducts, development of fibrosis, and granulomas, and may lead to cirrhosis [Webb et al., 2015]. The prevalence of PBC ranges from 1.91 to 40.2 per 100,000 people, with the important, yet unexplained, hallmark of predominance in females [Boonstra et al., 2012; Purohit et al., 2015]. Unfortunately, the factors leading to PBC are not fully understood. Environmental stimuli, as well as interacting immunogenetic and epigenetic risk factors, have been proposed contributors [Juran et al., 2014; Bianchi et al., 2014]. For example, pollution, tobacco smoke, toxin exposure, or infectious agents (e.g., *E. coli*) are considered promoters of PBC development [Lleo et al., 2017; Purohit et al., 2015; Carey et al., 2015]. Additionally, it has been shown by many studies that family history and genetic predisposition are important factors in PBC risk and etiology. In monozygotic twins, the concordance rate is 63%, and for the first-degree relatives of PBC patients, concordance is 4% [Lleo et al., 2017; Purohit et al., 2015].

The main symptoms of PBC are not specific and include fatigue, skin hyperpigmentation, pruritus (itchy skin), hepatosplenomegaly, and xanthelasmas. Portal hypertension or jaundice may also occur in PBC. However, 20%–60% of PBC cases are diagnosed in asymptomatic patients based on the detection of antimitochondrial autoantibodies (AMA), which are associated with the elevation of IgM and cholestatic biochemistry, and then confirmed by the presence of specific bile duct pathology assessed via liver biopsy [Lleo et al., 2017]. The presence of AMA, a highly specific marker that underscores the autoimmune nature of this disease, is identified in about 95% of patients with PBC. AMA associated with PBC are targeted to 2-oxo-acid dehydrogenase complexes, which include pyruvate dehydrogenase (PDC-E2), branched chain 2-oxo-acid dehydrogenase (BCOADC-E2), and 2-oxo-glutaric acid dehydrogenase (OADC-E2) [Carey et al., 2015].

Although it is clear that the pathophysiology of PBC is linked to an intense autoimmune response directed against biliary epithelial cells, the mechanism of biliary destruction remains enigmatic. The immunopathology of PBC is consistent with the dysregulation of both adaptive and innate immunity, resulting in excessive activity of various inflammatory cells [Ma and Chen, 2019]. Several specific mechanisms have been postulated, including IL-12-JAK-STAT4 signaling, Th1 T cell polarization, CD80 or IFN-gamma-dependent pathways, as well as the involvement of liver-infiltrating autoreactive T cells (CD4+ T cells, CD8+ T cells, natural killer T cells, and B cells) [Webb et al., 2015].

The activation of inflammatory cells is controlled on transcriptional and posttranscriptional levels by Monocyte Chemoattractant Protein-Induced Protein 1 (MCPIP1, *alias* Regnase1), which is involved in negative regulation of inflammation through its endonuclease activity [Matsushita et al., 2009; Mizgalska et al., 2009] that shortens the half-life of selected pro-inflammatory cytokine transcripts (e.g., IL-1β, IL-6, IL-8) and mitigates their function [Mino et al., 2015; Lipert et al., 2017; Dobosz et al., 2016]. The anti-inflammatory properties of MCPIP1 have been confirmed *in vivo* [Matsushita et al., 2009; Liang et al., 2010; Uehata et al., 2013; Li et al., 2017], but it is not known whether MCPIP1 controls the autoimmune response in patients with PBC. Recently, Sun et al. described a protective role of Mcpip1 in liver recovery after ischemia/reperfusion (I/R) injury, indicating that Mcpip1 functions to ameliorate liver damage, reduce inflammation, prevent cell death, and promote tissue regeneration [Sun et al., 2018], but did not study the role of MPCIP1 in the context of autoimmune liver disease.

To verify the hypothesis that Mcpip1 expression protects against the development of autoimmune liver disease, we generated mice lacking Mcpip1 in liver cells (Mcpip1^AlbKO^); their phenotype revealed increased levels of immunoglobulins (IgG, IgM) as well as AMA and anti-nuclear autoantibodies (ANA) autoantibodies (anti-PDC-E2, anti-gp-210), liver fibrosis, inflammation, and destruction of intrahepatic bile ducts that led to cholestasis. The phenotype of Mcpip1^AlbKO^ mice recapitulated features of human biliary cholangitis.

## Materials and methods

### Animals and genotyping

Animals were housed in ventilated cages under specific pathogen-free conditions in a temperature-controlled environment with a 14/10◻h light/dark cycle and fed *ad libitum*. To obtain hepatocyte-specific knockout of the *Zc3h12a* gene encoding Mcpip1, *Zc3h12a*^lox/lox^ mice (Mcpip1^fl/fl^), with loxp sites flanking exon 3 of the *Zc3h12a* gene, [Li et al., 2017], were crossed with liver-specific Cre-expressing transgenic mice (Alb^Cre tg/+^, Jackson Laboratory). The mice were designated Mcpip1^AlbKO^. Deletion of Mcpip1 in leukocytes of myeloid origin was achieved after crossing Mcpip1^fl/fl^ mice with a LysM-Cre expressing strain (LysM^Cre tg/+^, Jackson Laboratory). The mice were designated Mcpip1^LysMKO^. For genotyping, DNA was extracted from tail tissue using the KAPA Mouse Genotyping Kit (KAPA Biosystems) according to the manufacturer’s instructions. Genotyping for loxp insertion was performed by PCR using the following primers: GCCTTCCTGATCCTATTGGAG (wild-type), GAGATGGCGCAGCGCAATTAAT (knock-out), and GCCTCTTGTCACTCCCTCCTCC (common). Genotyping for Alb^Cre tg/+^ the following primers: TGCAAACATCACATGCACAC (wild-type), GAAGCAGAAGCTTAGGAAGATGG (mutant), and TTGGCCCCTTACCATAACTG (common), and genotyping for LysM^Cre^ was performed using TTACAGTCGGCCAGGCTGAC (wild-type), CCCAGAAATGCCAGATTACG (mutant), and CTTGGGCTGCCAGAATTTCTC (common). The Local Animal Ethical Commission approved all experiments.

### Histological evaluation of liver pathology in Mcpip1^AlbKO^ mice

Liver samples obtained from male and female mice of different ages were fixed in 4% buffered formalin and processed through the standard paraffin embedding method. Sections of 5 μm were stained with hematoxylin and eosin (HE) for general histology and Picro Sirius Red (PSR) for collagen deposition, and then visualized using a standard light microscope (Olympus BX51; Olympus Corporation). Qualitative evaluation of liver samples was performed under 200x and 400x magnification. For quantitative analysis of fibrosis, PSR-stained sections were imaged under 100x magnification and the degree of collagen deposition was analyzed using the Ilastik segmentation toolkit and Fiji ImageJ software (NIH, Bethesda, MD, United States). The number of bile ducts was calculated from counts within 10 randomly chosen portal areas per liver section, under 200x magnification in a blinded fashion, using a standard light microscope (Olympus BX41; Olympus Corporation).

### Transcriptome sequencing

Prior to library preparation, RNA sample integrity was evaluated and concentration was measured using an Agilent 2100 Bioanalyzer with an RNA 6000 Pico Kit (Agilent). Libraries for 16 samples were prepared using the Ion AmpliSeq™ Transcriptome Mouse Gene Expression Kit, which covers over 20,000 mouse RefSeq genes in a single assay. Libraries were prepared according to the manufacturer’s protocol using 50 ng of total RNA as input. Sixteen barcoded libraries were combined and pooled in equimolar concentrations and subsequently sequenced on two chips from the Ion PI Chip Kit v3 using an Ion Proton Sequencer and Ion PI Hi-Q Sequencing 200 chemistry. The primary bioinformatic analyses, including filtering and alignment of reads, were carried out using Torrent Suite Server v5.10.0. Subsequently, reads were counted and subjected to differential expression analysis using the DESeq2 package (with default parameters) implemented in R version 3.3.3. P-values for di[erentially expressed genes were corrected for multiple comparisons using the Benjamini-Hochberg approach and results with corrected p-values < 0.05 were considered significant. Functional annotation of di[erentially expressed genes (≥ 1.2-fold change) was performed using bioinformatic tools available via the Database for Annotation, Visualization and Integrated Discovery (DAVID; version 6.8; https://david.ncifcrf.gov/home.jsp). Gene lists were searched using Ensembl gene annotation (ENSEMBL_GENE_ID), with the *Mus musculus* background dataset used for analyses. Selected genes were mapped to Biological Process (BP) Gene Ontology (GO) terms. Venn diagrams were created using the InteractiVenn tool [Heberle et al., 2015]. Principal component analysis (PCA) was conducted for RNA-Seq data to demonstrate differential gene expression in the analyzed cell types. PCA analysis and subsequent plot generation were conducted using DESeq2 and ggplot2 libraries in R.

### Statistical analysis

Results are expressed as mean ± SEM. The exact number of experiments or animals used is indicated in figure legends. Two-tailed Student’s t-test was used for comparison of two groups, and one-way ANOVA with Bonferroni post hoc test was applied for comparison of multiple groups. Statistical significance is indicated by asterisks in figures, with the following definitions: * p<0.05; ** p<0.01; *** p<0.001.

For further details regarding the materials and methods used, please refer to the supplementary information.

## Results

### Development of liver-specific *Zc3h12a-*deficient mice

To study the function of Mcpip1 protein *in vivo*, we generated Mcpip1^AlbKO^ knockout mice which lack the *Zc3h12a* gene in liver cells. As previously described by Li and colleagues [Li et al., 2017], murine embryonic stem cells were genetically modified to obtain a loxp knock-in flanking the 3^rd^ exon of the *Zc3h12a* gene. After generation of chimeric mice and a series of backcrosses, the C57BL/6N *Zc3h12a* fl/fl mice were obtained (Fig. S1A). Control Mcpip1^fl/fl^ and Mcpip1^AlbKO^ mice were selected after standard genotyping (Fig. S1B), and deletion of the *Zc3h12a* gene was confirmed in Mcpip1^AlbKO^ mice at both the RNA and protein levels (Fig. S1C,D). The knockout mice showed diminished *Zc3h12a* expression in the liver and primary hepatocytes, but not in the heart and lungs, confirming liver specificity of Mcpip1 silencing (Fig. S1C,D).

### General characteristics of Mcpip1^AlbKO^ mice

Mcpip1^AlbKO^ mice were born at the expected Mendelian ratio and had no gross abnormalities or impairment in general fitness up to 52 weeks of age compared to Mcpip^fl/fl^ mice. There were no differences in body mass of Mcpip1^AlbKO^ versus control mice, however, Mcpip1^AlbKO^ animals were characterized by hepatomegaly and splenomegaly (Fig. S1E-G). Blood morphology parameters of 6, 24, and 52-week-old Mcpip1^AlbKO^ animals did not differ significantly from Mcpip1^fl/fl^ counterparts (Table 1). Alkaline phosphatase (ALP) activity and concentration of total bile acids (TBA) in the serum of 6-week-old Mcpip1^AlbKO^ mice were both significantly higher than in control animals (Table 2). In 52-week-old Mcpip1^AlbKO^ mice, TBA was also higher compared with that of age-matched controls, but the difference did not reach significance (p=0.083). In contrast to young and old Mcpip1^AlbKO^ mice, 24-week-old Mcpip1^AlbKO^ mice did not show any differences in biochemical parameters compared with their Mcpip1^fl/fl^ counterparts (Table 2).

**Table 1.** Blood counts of Mcpip1^fl/fl^ and Mcpip1^AlbKO^ mice.

| Component | 6 weeks |  | 24 weeks |  | 52 weeks |  |
| --- | --- | --- | --- | --- | --- | --- |
|  | Mcpip1<sup>fl/fl</sup> | Mcpip1<sup>AlbKO</sup> | Mcpip1<sup>fl/fl</sup> | Mcpip1<sup>AlbKO</sup> | Mcpip1<sup>fl/fl</sup> | Mcpip1<sup>AlbKO</sup> |
| Erythrocytes (*10<sup>12</sup>/L) | 9.19 ± 0.62 | 8.98 ± 0.83 | 9.73 ± 0.85 | 9.28 ± 0.61 | 10.45 ± 0.23 | 9.76 ± 0.26 |
| Hemoglobin (g/dL) | 15.03 ± 0.63 | 14.83 ± 0.98 | 14.52 ± 1.10 | 14.13 ± 0.93 | 14.75 ± 0.32 | 14.14 ± 0.31 |
| Hematocrit (%) | 46.19 ± 2.64 | 45.68 ± 4.28 | 45.67 ± 4.76 | 44.01 ± 2.57 | 51.17 ± 1.29 | 48.70 ± 1.17 |
| MCV (fl) | 50.43 ± 1.27 | 51.00 ± 0.82 | 46.83 ± 1.47 | 47.89 ± 1.17 | 49.17 ± 0.70 | 49.78 ± 0.57 |
| MCH (pg) | 16.39 ± 0.48 | 16.60 ± 0.42 | 14.93 ± 0.42 | 15.28 ± 0.19 | 14.12 ± 0.21 | 14.50 ± 0.20 |
| MCHC (g/dL) | 32.59 ± 0.81 | 32.53 ± 1.07 | 31.85 ± 0.87 | 31.89 ± 0.51 | 28.82 ± 0.27 | 29.03 ± 0.19 |
| Platelets (*10<sup>9</sup>/L) | 749.0 ± 125.8 | 672.8 ± 80.1 | 989.3 ± 192.2 | 1051.3 ± 248.1 | 1314.6 ± 91.8 | 1342.3 ± 192.4 |
| Leukocytes (*10<sup>9</sup>/L) | 3.74 ± 0.53 | 5.17 ± 0.56 | 5.97 ± 0.66 | 6.46 ± 0.65 | 5.65 ± 0.31 | 6.64 ± 0.82 |
| Lymphocytes (*10<sup>9</sup>/L) | 2.09 ± 0.30 | 3.33 ± 0.29 | 3.90 ± 0.86 | 3.87 ± 1.16 | 3.88 ± 0.34 | 4.84 ± 1.07 |
| Granulocytes (*10<sup>9</sup>/L) | 1.51 ± 0.20 | 1.48 ± 0.28 | 2.53 ± 0.56 | 2.37 ± 0.38 | 1.83 ± 0.11 | 2.74 ± 1.48 |
| Monocytes (*10<sup>9</sup>/L) | 0.14 ± 0.04 | 0.28 ± 0.03 | 0.25 ± 0.04 | 0.31 ± 0.08 | 0.30 ± 0.04 | 0.40 ± 0.24 |

Mean ± SEM, for Mcpip1<sup>fl/fl</sup> n=5 and for Mcpip1<sup>AlbKO</sup> n=8.

**Table 2.** Blood biochemistry of Mcpip1^fl/fl^ and Mcpip1^AlbKO^ mice.

| Component | 6 weeks |  | 24 weeks |  | 52 weeks |  |
| --- | --- | --- | --- | --- | --- | --- |
|  | Mcpip1<sup>fl/fl</sup> | Mcpip1<sup>AlbKO</sup> | Mcpip1<sup>fl/fl</sup> | Mcpip1<sup>AlbKO</sup> | Mcpip1<sup>fl/fl</sup> | Mcpip1<sup>AlbKO</sup> |
| ALT (U/L) | 46.60 ± 6.91 | 50.36 ± 2.57 | 53.64 ± 8.28 | 48.39 ± 7.59 | 38.14 ± 6.37 | 92.40 ± 33.58 |
| Albumin (g/dL) | 3.33 ± 0.07 | 3.36 ± 0.05 | 3.15 ± 0.11 | 3.19 ± 0.09 | 3.64 ± 0.07 | 3.46 ± 0.13 |
| AST (U/L) | 125.9 ± 15.7 | 119.5 ± 12.6 | 141.8 ± 17.4 | 152.5 ± 10.8 | 116.5 ± 16.6 | 180.3 ± 46.9 |
| Bilirubin (mg/dL) | 0.06 ± 0.01 | 0.05 ± 0.01 | 0.10 ± 0.03 | 0.08 ± 0.01 | Not detected | Not detected |
| Cholesterol (mg/dL) | 99.11 ± 6.74 | 111.10 ± 3.01 | 95.14 ± 7.15 | 105.40 ± 5.73 | 93.25 ± 8.52 | 92.00 ± 9.63 |
| ALP (U/L) | **149.0 ± 12.0** | **287.7 ± 43.2 \*\*** | 72.4 ± 1.8 | 75.1 ± 4.5 | 83.1 ± 4.6 | 108.3 ± 21.8 |
| TBA (μmol/L) | **7.3 ± 3.1** | **111.7 ± 28.2 \*\*** | Not analyzed | Not analyzed | 40.9 ± 23.6 | 90.1 ± 21.6 <sup>p=0.083</sup> |

Mean ± SEM, for Mcpip1<sup>fl/fl</sup> n=5 and for Mcpip1<sup>AlbKO</sup> n=8.

### Histological and serological pathology of Mcpip1^AlbKO^ mice recapitulates human primary biliary cholangitis

Mcpip1^AlbKO^ mice at 6-52 weeks of age, but not Mcpip1^fl/fl^ mice, displayed characteristic liver pathology, including active and progressive proliferation of intrahepatic bile ducts accompanied by extensive parenchymal inflammation and fibrosis, along with fibrosis in portal areas, all at various intensities depending on age (Fig. 1, S2, S3). In general, all features of intrahepatic bile duct pathology were extensive in 6-week-old Mcpip1^AlbKO^ mice, however, at 24 weeks of age, histopathological changes were less severe in the liver parenchyma. By 52 weeks of age, the histopathological features of PBC were re-intensified in Mcpip1^AlbKO^ mice (Fig. 1C,D,G,H).

**Fig. 1.**
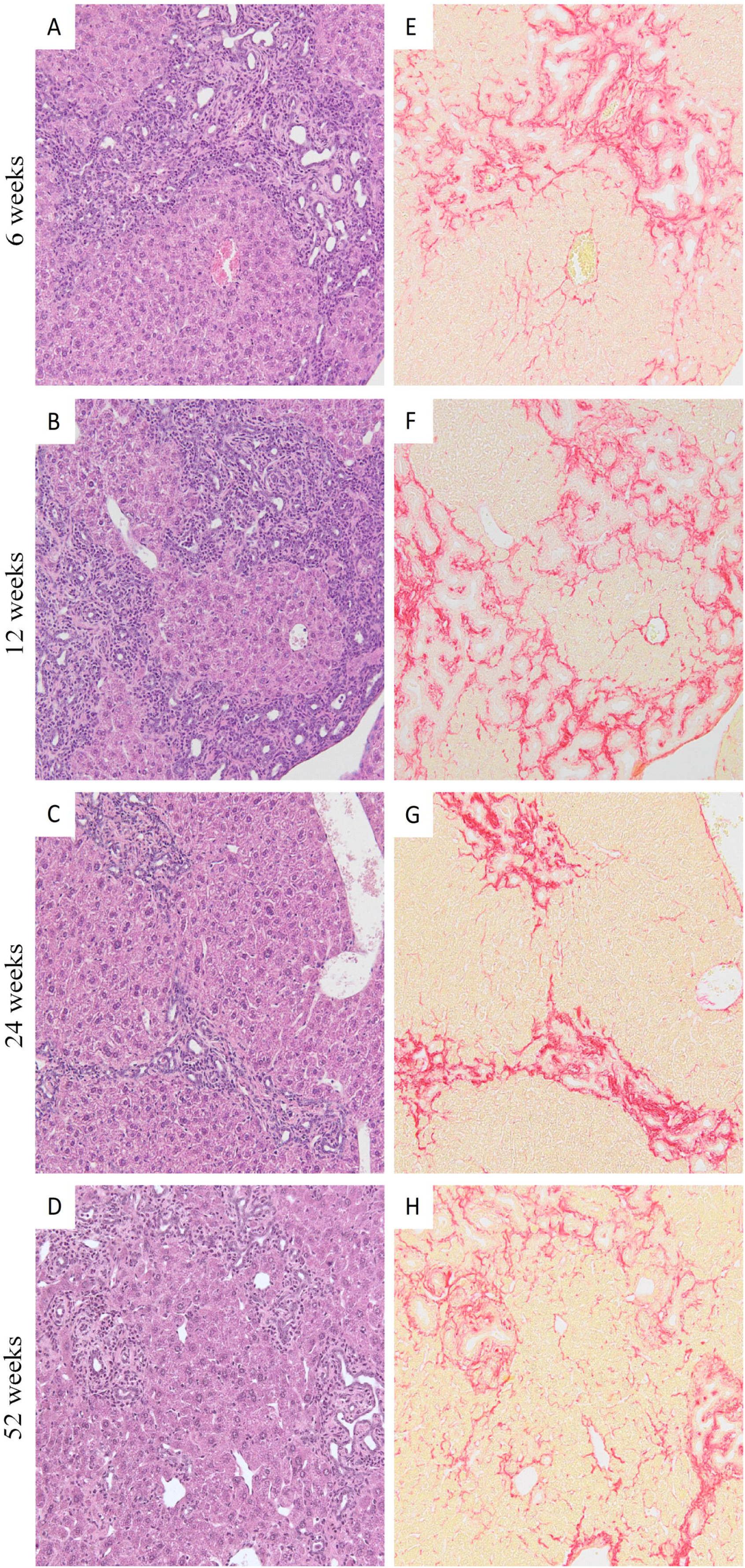
Progression of intrahepatic bile duct pathology in the liver parenchyma of 6 to 52-week-old Mcpip1^AlbKO^ male mice. Proliferative changes in intrahepatic bile ducts with inflammatory infiltration predominate in young (6-week-old) mice leading to bile duct destruction, intraductal obstruction, and extensive fibrosis. At the age of 24 weeks, the processes of proliferation, inflammation, and collagen deposition were reduced, while aging mice (52-week-old) developed new foci of cholangiocyte proliferation accompanied by active fibrotic areas. Magnification 200x, HE (A-D) and PSR (E-H) staining. HE, hematoxylin and eosin; PSR, picrosirius red.

In 6-week-old Mcpip1^AlbKO^ mice, active and intensive collagen deposition was associated with extensive bile duct hyperplasia and inflammation (Fig. 1E, S3). At the ages of 12 and 24 weeks, proliferation and inflammatory processes diminished temporarily and the area of fibrotic tissue was reduced (Fig. 1F,G, S3). However, recurrence of intrahepatic and periportal proliferation with inflammatory infiltration in livers of 52-week old animals led to reactivation of collagen deposition (Fig. 1H, S3). Unlike younger animals, 52-week-old Mcpip1^AlbKO^ mice showed that proliferative processes were not limited to intrahepatic bile ducts but active along sinusoids throughout the parenchyma (Fig. 1D). The entire spectrum of features of intrahepatic bile duct pathology that occurred during disease progression was present in 52-week-old Mcpip1^AlbKO^ mice, as shown in Fig. 2 (Fig. 2A-H). The extensive proliferation of bile ducts in the liver parenchyma and portal areas was accompanied by bile duct epithelium disruption, inflammation, and fibrosis, resulting in lumen obstruction and bile duct destruction (Fig. 2A-C). Eventually, the closure of intrahepatic bile duct lumen, bile acid deposition in hepatocytes, and formation of small granulomas were observed (Fig. 2D-F).

**Fig. 2.**
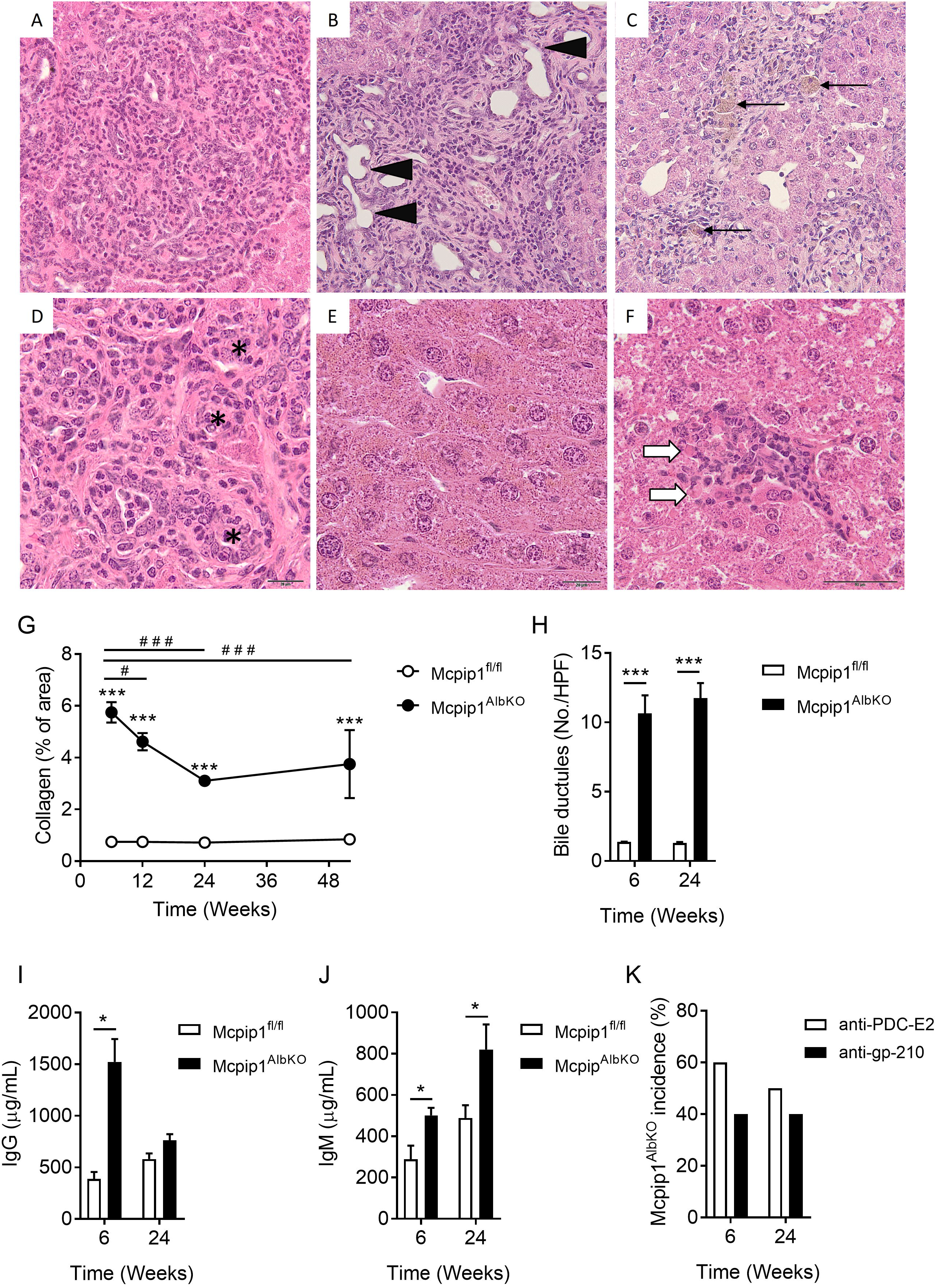
Characterization of hallmarks of primary biliary cholangitis in Mcpip1^AlbKO^ mice. HE staining of 52-week old mice showing (A) Proliferation of intrahepatic bile ducts with associated parenchymal inflammation; (B) Epithelial destruction of intrahepatic bile ducts (black arrows) and extensive fibrosis of parenchyma; (C) Resolution of inflammation in intrahepatic bile ducts involving macrophages (black arrows); (D) Dysplasia of cholangiocytes with associated intraductal obstruction of intrahepatic bile ducts (asterisks); (E) Cholestasis and bile acid deposition in hepatocytes; (F) Small granuloma (white arrows indicate hepatocyte necrosis), magnification 400x. (G) Collagen area in the liver of 6, 12, 24, and 52-week-old Mcpip1^fl/fl^ and Mcpip1^AlbKO^ mice; (H) Number of bile ductules in livers per high power field (HPF); Concentrations of total IgG (I) and IgM (J) classes in plasma of 6 and 24-week-old Mcpip1^fl/fl^ and Mcpip1^AlbKO^ mice; (K) Percentage of Mcpip1^AlbKO^ mice, with higher concentration of anti-PDC-E2 antimitochondrial autoantibodies and anti-gp-210 antinuclear autoantibodies compared to mean concentration in control Mcpip1^fl/fl^ counterparts. Data represent mean ± SEM, \**p*◻<◻0.05, \*\*\**p* < 0.001 vs. Mcpip1 fl/fl; #*p* < 0.05, ##*p* < 0.01; ###*p* < 0.001 vs. 6 weeks. H&E, hematoxylin and eosin.

To further assess the similarities between the phenotype of Mcpip1^AlbKO^ mice and the characteristics of PBC in humans, levels of total IgG, IgM, and specific AMA and ANA were measured. IgG and IgM concentrations were higher in 6 and 24-week-old Mcpip1^AlbKO^ animals compared to Mcpip1^fl/fl^ counterparts (Fig. 2I,J). Additionally, 50–60% of Mcpip1^AlbKO^ mice had higher plasma concentrations of anti-PDC-E2 mitochondrial autoantibodies than control counterparts (Fig. 2K). Similarly, incidence of anti-gp-210 nuclear autoantibodies, was approximately 40% in Mcpip1^AlbKO^ mice (Fig. 2K).

To verify whether systemic inflammation caused by lack of Mcpip1 protein in myeloid cells might also lead to PBC development, Mcpip1^LysMKO^ mice were analyzed. Features of human PBC were not observed in Mcpip1^LysMKO^ mice, despite the presence of robust myeloid cell-driven systemic inflammation (Fig. S4).

### Male and female Mcpip1AlbKO mice have the same pathological features

In humans, PBC is disease with high predominance in females. We therefore compared the phenotype of female and male Mcpip1^AlbKO^ mice. Histological comparison of liver sections from 6, 12, and 24-week-old Mcpip1^AlbKO^ mice revealed no differences in the development of pathology between female and male mice. All characteristic features of intrahepatic bile duct pathology demonstrated in male Mcpip1^AlbKO^ mice were also present in female Mcpip1^AlbKO^ mice, including a similar pattern of collagen deposition in the liver (Fig. S5A). There was also no difference between male and female Mcpip1^AlbKO^ mice in plasma levels of anti-PDC-E2 and anti-gp-210 autoantibodies (Fig. S5B).

### Transcriptome changes in Mcpip1-depleted primary hepatocytes

To better understand the molecular mechanisms responsible for the robust PBC phenotype in 6-week-old Mcpip1^AlbKO^ mice and its downregulation in 24-week-old Mcpip1^AlbKO^ mice, the transcriptome of isolated primary hepatocytes was analyzed. Differential gene expression analysis followed by calculation of p-values revealed that, in 6 and 24-week-old Mcpip1^AlbKO^ mice, there were 812 and 8 differentially expressed genes (DEGs), respectively, compared with their respective age-matched controls.

In hepatocytes derived from 6-week-old Mcpip1^AlbKO^ mice, 688 DEGs were upregulated and 124 were downregulated (Table S1 in the Supplementary Material). As shown in Figure 3, upregulated genes were related to cell cycle control (e.g., cell division, mitotic nuclear division, cell cycle, positive regulation of cell proliferation), regulation of inflammation (e.g., inflammatory response, immune response, cytokine-mediated signaling, IL-1β and IL-6 production), and extracellular matrix biosynthesis (e.g., extracellular matrix and collagen fibril organization, collagen biosynthesis, extracellular matrix constituent secretion). Downregulated genes were related to lipid metabolism (e.g., lipid metabolic processes, fatty acid (FA) β-oxidation, FA and cholesterol metabolic processes, FA transport, negative regulation of lipid storage), oxidation-reduction processes, glucose homeostasis, and organic anion transport (Fig. 3).

**Fig. 3.**
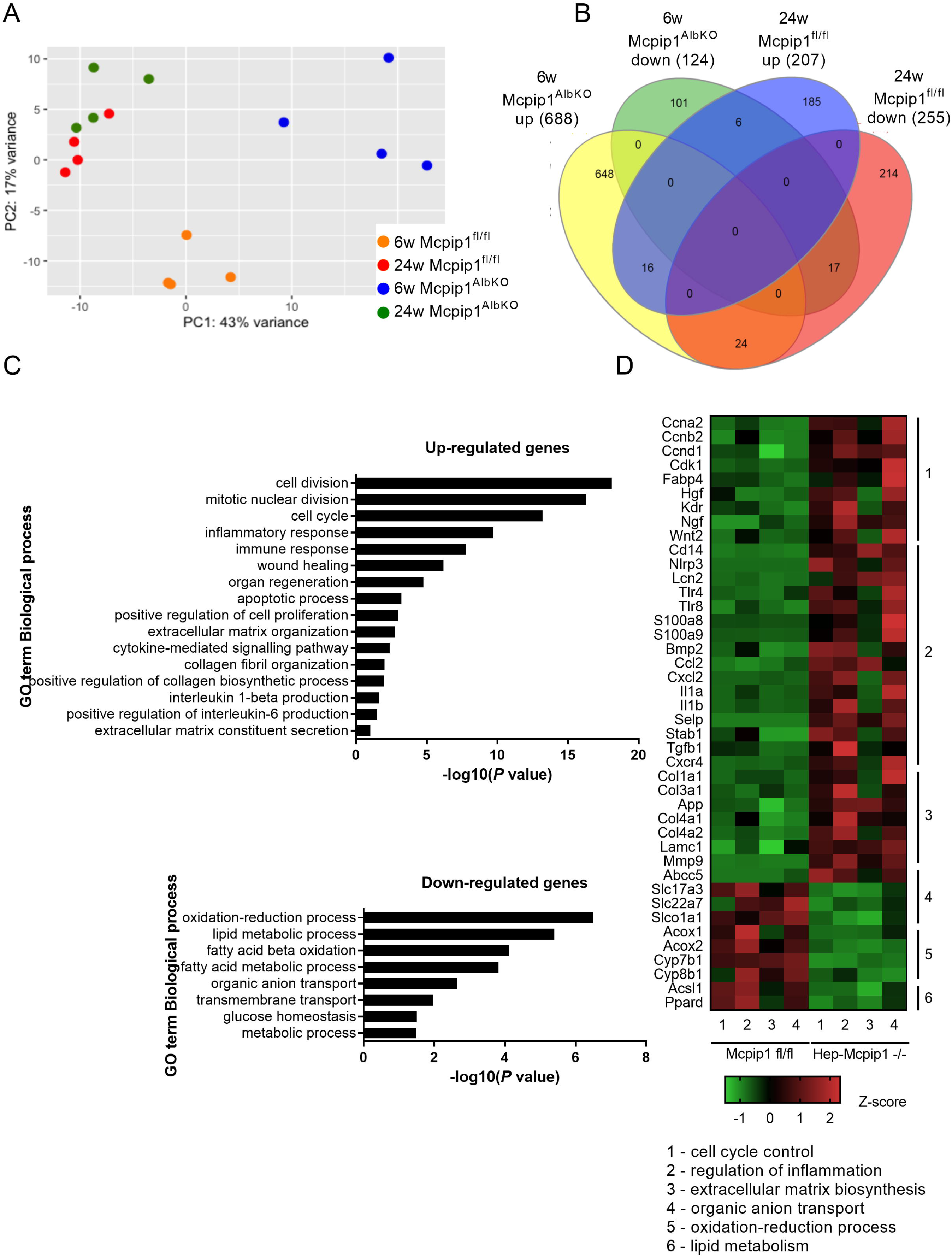
Transcriptome analysis of Mcpip1^fl/fl^ and Mcpip1^AlbKO^ primary hepatocytes. (A) PCA plot for RNA-Seq dataset demonstrating differential gene expression in all analyzed primary hepatocytes isolated from 6 and 24-week-old Mcpip1^fl/fl^ (6w Mcpip1^fl/fl^, 24w Mcpip1^fl/fl^, respectively) and 6 and 24-week-old Mcpip1^AlbKO^ mice (6w Mcpip1^AlbKO^, 24w Mcpip1^AlbKO^, respectively); (B) Venn diagram presenting the number of transcripts significantly up- and downregulated (adj. P-value < 0.05) in 6-week-old Mcpip1^AlbKO^ samples (6w Mcpip1^AlbKO^ up/down) and 24-week-old Mcpip1^fl/fl^ (24w Mcpip1^fl/fl^ up/down) in comparison to 6-week-old Mcpip1^fl/fl^ controls (6w Mcpip1^fl/fl^); (C) GO enrichment analysis of genes differentially expressed in 6-week-old Mcpip1^fl/fl^ and Mcpip1^AlbKO^ primary hepatocytes. Scale is the −log10(P-value) of the enrichment score (p-value < 0.05); (D) Heat map showing differentially expressed genes selected from GO enrichment analysis.

In hepatocytes from 24-week-old Mcpip1^AlbKO^ mice, only 8 genes were differently expressed compared to age-matched control mice (upregulated in Mcpip1^AlbKO^: *Lpin1, Lrp6,* and *Tmc7*; downregulated in Mcpip1^AlbKO^: *Cela1, Ppdpf, Asap1, Lect2,* and *Pla2g16*; Table S2 in the Supplementary Material).

The striking difference between the transcriptomes of 6 and 24-week-old Mcpip1^AlbKO^ versus their respective controls was also confirmed by direct comparison of the transcriptomes of hepatocytes from 6 and 24-week-old Mcpip1^AlbKO^ mice, which showed 725 DEGs, with 538 upregulated and 187 downregulated (Fig. S5, Table S3 in the Supplementary Material). This comparison revealed similar enhanced processes (mitotic nuclear division, cell cycle, and cell division) when hepatocytes isolated from 6-week-old control and knockout mice were compared.

## Discussion

We have, to our knowledge, been the first to demonstrate that the phenotype of Mcpip1^AlbKO^ mice recapitulates most features of human PBC (Table 3). On a histopathological level, Mcpip1^AlbKO^ mice displayed intrahepatic bile duct pathology that included active proliferative changes, inflammatory infiltration, bile duct epithelium disruption, small granuloma formation, and significant and progressive liver fibrosis in the parenchyma and portal areas. On a functional level, Mcpip1^AlbKO^ mice showed cholestasis. Autoimmune response, a characteristic feature of PBC, was also detected in Mcpip1^AlbKO^ mice, as demonstrated by increased plasma concentrations of IgG and IgM, as well as AMA and ANA autoantibodies (anti-PDC-E2, anti-gp-210). As Mcpip1 deficiency in myeloid cells (Mcpip1^LysMKO^ mice) did not result in features of PBC, we concluded that Mcpip1, specifically in the liver, is a negative regulator of autoimmune response and may protect against PBC development. Importantly, our results showed that Mcpip1^AlbKO^ mice had robust postnatal symptomatology suggestive of prenatal origin, displayed a mild phenotype in middle age, and regained robust PBC features with advanced age. Transcriptomic analysis of primary hepatocytes revealed a distinct set of genes compared to those previously linked to PBC, and identified distinct transcriptomes for active and chronic phases of PBC. Altogether, Mcpip1^AlbKO^ mice represent a unique model that closely mimics PBC in humans (see Table 3 for summary) and has significant potential to increase insight into the pathophysiology of PBC.

**Table 3.** Comparison between characteristics of human PBC and Mcpip1^AlbKO^ mice.

| Feature | Human PBC | Mcpip1<sup>AlbKO</sup> |
| --- | --- | --- |
| Classification | Complex etiology | Genetic |
| Gender differences | Female predominance | No |
| Environmental factor | Yes | No |
| Elevated serum bile acids | Yes | Yes |
| Presence of autoantibodies: |  |  |
| AMA | 90-95% | 50-60% |
| ANA | 40-50% | 40% |
| Immunoglobulins | IgM +; IgG + | IgM +; IgG + |
| Liver histology: |  |  |
| Lymphocytic infiltration | Yes | Yes |
| Bile duct destruction | Yes | Yes |
| Granuloma | Yes | Yes |
| Fibrosis | Yes | Yes |
| Other findings |  | PBC phenotype was robust in 6-week-old mice, diminished in 24-week-old mice and reintensified in 52-week-old mice |

Previously described murine models of PBC, including genetically modified models, xenobiotic immunized models, and infection-triggered models display some important features of PBC, such as the presence of AMA, elevated levels of IgG and IgM, portal lymphocyte infiltration, and bile duct destruction [Katsumi et al., 2015]. However, they do not recapitulate all features of PBC; for example, other mouse models do not display progressive fibrosis, a central clinical feature that informs prognosis, including morbidity and mortality, in human PBC patients [Katsumi et al., 2015]. Although Mcpip1^AlbKO^ mice do not show female predominance of PBC features, which has been observed in only one PBC model (IFNγ-ARE-Del mice) [Bae et al., 2016], the Mcpip1^AlbKO^ mouse model has the advantage of clinical relevance via its unambiguous histopathological features and full spectrum of PBC-associated pathological changes that are not comprehensively present in any previously described model (fig 2A-H, Fig. 1, S2, S3). Importantly, fibrosis was present in 100% of Mcpip1^AlbKO^ mice, compared to approximately 50% of IL-12p35−/−;dnTGFβRII double mutant mice [Tsuda et al., 2013]. Another unique feature of Mcpip1^AlbKO^ mice is the robust postnatal PBC phenotype followed by remission in middle age, and age-dependent disease progression that is not driven by systemic inflammation, but by intrinsic liver-derived pathology. Although it is not known whether PBC in humans has prenatal origin and MCPIP1-driven pathogenesis, peak incidence of PBC in humans occurs in the fifth decade of life; timing that is compatible with the mild phenotype in middle-aged Mcpip1^AlbKO^ mice followed by age-dependent progression of the disease.

The Mcpip1^AlbKO^ mouse model of PBC presented here differs from other models on both the phenotype and genomic levels. Previously, PBC mouse model development was driven by alteration of function of various genes, including *Ifng* [Bae et al., 2016], *Tgfbr2* [Oertelt et al., 2006], *Il2ra* (IL-2Rα−/−) [Wakabayashi et al., 2006], *Foxp3* [Zhang et al., 2009], *Il12a* [Tsuda et al., 2013] and *Slc4a2* [Salas et al., 2008], yet none of these genes was found to be differentially expressed in hepatocytes isolated from Mcpip1^AlbKO^ mice. Indeed, Mcpip1^AlbKO^ mice displayed a unique set of DEGs in hepatocytes, with a strikingly distinct profile of DEGs in active and chronic phases of the disease in 6 and 24-week-old Mcpip1^AlbKO^ mice, respectively.

In 6-week-old Mcpip1^AlbKO^ mice with acute phase PBC, we identified enhanced expression of many genes involved in cell proliferation (e.g., *Hgf, Tgfb1, Wnt2*), as well as genes involved in biliary development and cholangiocyte proliferation and important for biliary repair (e.g., *Hgf, Pdgfa, Tgfb1, Bmp2, Mmp9*), fibrosis (e.g., *Tgfb1, Lpin1*), inflammation (e.g., *Il1a, Il1b, ccl2, Cxcl2, Cxcl12, S100a8, S100a9*), or extracellular matrix organization (e.g., *Mmp9, Col1a1, Col3a1*).

In contrast to acute phase PBC with 812 DEGs, the chronic remission phase showed only 8 DEGs, including altered expression of *Lrp6*, *Lect2*, and *Lipin-1*. Low-density lipoprotein receptor-related protein 6 (Lrp6) linked with the Wnt/β-catenin signaling pathway was shown to regulate liver fibrosis [Yu et al., 2020]. Leukocyte cell-derived chemotaxin-2 (*LECT2*), originally identified as a hepatocyte-secreted chemokine [Lebensztejn et al., 20116] was identified as a prognostic indicator in acute liver failure [Sato et al., 2004; Slowik et al., 2019]. Finally, *Lipin-1* (*Lpin1*), the only gene with significantly different expression in knockout cells at both time points, was shown to coactivate HNF4α and regulate fatty acid catabolism [Chen et al., 2012] and TGF-β-induced fibrogenesis [Jang et al., 2016].

Mcpip1^AlbKO^ mice also showed altered expression of organic anion transporters encoding Organic anion transporting polypeptide 1a1 (Oatp1a1 protein, *Slco1a1* gene) and Organic anion transporter 2 (Oat2 protein, *Slc22a7* gene), which are involved in bile acid transport [Roth et al., 2012]. These changes likely secondary to the PBC phenotype in Mcpip1^AlbKO^ mice and compatible with the changes in expression of other organic transporters, such as NTCP, OATP1B1, and OATP1B3 in humans with PBC [Thakkar et al., 2017].

Although the expression of multiple genes were altered by the deletion of Mcpip1 alone in the acute phase of PBC, it seems impossible to identify a single signaling pathway responsible for the observed PBC phenotype in Mcpip1^AlbKO^ mice, and changes in expression of multiple genes and their interactions are likely responsible. Although some genes differently expressed in the remission phase could be linked to liver fibrosis and liver failure, their roles in this PBC phenotype remain to be established, along with links between the acute and chronic phases of PBC and the mechanisms of activation of the disease phenotype in aging Mcpip1^AlbKO^ mice.

Nevertheless, our work has demonstrated the novel role of Mcpip1 as a gene orchestrating biliary duct hemostasis and identified that the PBC phenotype results from Mcpip1 deficiency in the liver. Thus far, MCPIP1 is primarily known for its anti-inflammatory properties that are mediated *via* endonuclease activity and result in shortening the half-life of selected pro-inflammatory cytokine transcripts (e.g., IL-1β, IL-6, IL-8) and mitigating their function [Matsushita et al., 2009; Mizgalska et al., 2009; Mino et al., 2015; Dobosz et al., 2016]. In a number of reports, the role of Mcpip1 in immune response has been demonstrated *in vivo* [Matsushita et al., 2009; Liang et al., 2010; Yu et al., 2013; Peng et al., 2018] and some reports suggested the involvement of Mcpip1 in autoimmune response; for example, in autoimmune gastritis and lupus [Zhou et al., 2013; Koziel et al. submitted]. Our study extends the knowledge of Mcpip1 as a regulator of autoimmune response in the liver, as we demonstrated that Mcpip1 deficiency triggered autoimmunity against bile ducts and led to the PBC phenotype.

In conclusion, our study demonstrated that Mcpip1 expression in the liver is a key protective factor against development of PBC. Mcpip1^AlbKO^ mice represent an excellent model through which to investigate the molecular mechanisms responsible for PBC development and test novel treatment strategies for PBC.

## Abbreviations

ALP: alkaline phosphatase
AMA: antimitochondrial autoantibodies
ANA: anti-nuclear autoantibodies
BSA: bovine serum albumin
DEG: differentially expressed gene
FA: fatty acid
HE: hematoxylin and eosin
I/R: ischemia/reperfusion
LC-MS: liquid chromatography-mass spectrometry
LSEC: liver sinusoidal endothelial cells
PBC: primary biliary cholangitis
PBS: phosphate-buffered saline
PCA: principal component analysis
TBA: total bile acid

## Authors contributions

Concept and design: JeK, JoK, SCh; Data acquisition: JeK, TH, ICC, MW, NP, AJ, AK, ED, ML, KM; analysis of RNA-seq data: EP; Providing Mcpip1^fl/fl^ mice: MF; Analysis and interpretation of data: JeK, TH, AJ, JJ, JoK, SCh; Writing and correcting the manuscript: JeK, JJ, JoK, SCh,

