## Supplementary information for "Deletion of Mcpip1 in Mcpip1^AlbKO^ mice recapitulates the phenotype of human primary biliary cholangitis"

Jerzy Kotlinowski, Address: Gronostajowa Street 7, 30-387 Krakow, Poland.

\# equal contribution as a senior author

**Keywords**

Primary biliary cholangitis, MCPIP1, Regnase1.

**Supplementary Materials and methods**

Blood count and blood biochemistry

Blood was collected by cardiac puncture and full blood count was performed using SCIL Vet ABC Plus or hematology blood analyzer (HORIBA Medical, Montpellier, France). The biochemistry profile was assessed using the Cobas 6000 analyzer (Roche Diagnostics, Indianapolis, USA).

Measurement of cytokine concentrations

The plasma concentration of cytokines IL-6, IL-10, MCP-1, IFN-γ, TNF-α, and IL-12p70 was measured using the cytometric bead array (BD Biosciences), according to the manufacturer’s instructions.

Measurement of total immunoglobulins

Mouse ELISA Quantitation assays (Bethyl Laboratories) were used to evaluate levels of IgG and IgM immunoglobulins in murine plasma samples. After diluting samples, tests were performed according to the manufacturer’s instructions. Absorbance was measured at 450 nm using a microplate reader (Tecan). All experiments were performed in duplicate.

Measurement of anti-PDC-E2 antibody levels

To evaluate the level of anti-PDC-E2 autoantibodies in plasma samples, the ELISA assay was performed with triple washing with phosphate-buffered saline (PBS) between all incubations. Microtiter wells (Bethyl Laboratories) were coated with PDC-E2 recombinant protein (3 µg/mL; MyBioSource) and then blocked with 2% bovine serum albumin (BSA) in PBS. Consecutively, plasma samples (diluted 10x) and then anti-mouse IgG, IgM, or IgA (H+L) secondary antibodies (1:1000; Thermo Fisher Scientific) diluted in 0.5% BSA in PBS were added. Next, samples were incubated with TMB HRP Substrate (Bethyl Laboratories) for 15 minutes, followed by the addition of Stop Solution (Bethyl Laboratories). Absorbance was measured at 450 nm using a Tecan Spectra Fluor Plus Microplate Reader. Experiments were performed in duplicate. First, the mean concentration of anti-PDC-E2 autoantibodies in Mcpip1^fl/fl^ mice was calculated. Next, we analyzed the percentage of Mcpip1^AlbKO^ animals with a higher concentration of anti-PDC-E2 in comparison to the mean value obtained for corresponding Mcpip1^fl/fl^ mice; results from Mcpip1^AlbKO^ mice are presented as incidence (%).

Measurements of anti-nuclear antibodies

Commercially available ELISA assays (Creative Diagnostics) were performed to evaluate levels of gp-210 ANA in murine plasma samples. The test was conducted according to the manufacturer’s instructions. Absorbance was measured at 450 nm using microplate reader. Experiments were performed in duplicate and results were calculated and presented as incidence (%).

Primary liver cell isolation

Primary liver sinusoidal endothelial cells (LSEC) were isolated from 24-week old male Mcpip1^fl/fl^, Mcpip1^AlbKO^, and Mcpip1^LysMKO^ mice according to the protocol described by Kus et al. (2019). After isolation, LSEC were incubated overnight in EBM-2 Basal Medium (Lonza) supplemented with 1% Antibiotic Antimycotic Solution for Cell Culture (Sigma-Aldrich), at 37°C and 5% CO_2_. Next, medium samples were collected for prostanoid analysis by liquid chromatography-mass spectrometry (LC-MS/MS).

Primary hepatocytes were obtained from Mcpip1^fl/fl^ and Mcpip1^AlbKO^ mice *via* collagenase perfusion, as described previously [Pydyn et al., 2019]. Briefly, animals were anesthetized with intraperitoneal administration of ketamine (100 mg/kg) and xylazine (10 mg/kg). Next, livers were perfused, *via* the inferior vena cava, with 20 mL Krebs-Ringer buffer supplemented with 0.1 mM EDTA followed by 30 mL of digestion solution (Krebs-Ringer buffer supplemented with 4.76 mM CaCl_2_ and 200 U/mL collagenase IV (Gibco). After excising the liver, it was disrupted in a Petri dish containing 10 mL of complete medium and filtered through a 100 µm cell strainer. The flow-through was centrifuged (500 *g*, 3 min, 4ºC) and the pellet was suspended in 10 mL of culture medium and then carefully layered on a Percoll gradient (4 mL of 1.12 g/mL, 5 mL of 1.08 g/mL and 5 mL of 1.06 g/mL). After Percoll gradient centrifugation, the two upper layers containing cell debris and non-parenchymal cells were carefully pipetted out and discarded. The lowest layer, which contained hepatocytes, was collected and washed. The viability of the isolated hepatocytes, estimated by trypan blue staining, was usually 80–90%. Directly after isolation, the primary hepatocytes were lysed in RIPA buffer for protein isolation. Total RNA was isolated with mirVana according to the manufacturer’s instructions (Thermofisher, Waltham, United States).

Quantification of prostanoid release by LSEC

The quantification of selected prostaglandins (6-keto-PGF1α, TBX_2_, and PGD_2_) in culture medium samples was performed using a UFLC Nexera liquid chromatography system (Shimadzu, Kyoto, Japan) coupled to a QTrap 5500 triple quadrupole mass spectrometer (Sciex, Framingham, MA, United States) according to the methodology described previously [Kij et al., 2020].

RNA isolation and RT-PCR

Total RNA and protein isolated from bone marrow from femur and tibia, as well as hearts, livers, lungs, and spleens were used to confirm tissue-specific deletion of Mcpip1. Total RNA was isolated using Fenozol (A&A Biotechnology). The concentration of total RNA was assessed using a NanoDrop 1000 Spectrophotometer (Thermo Fisher Scientiﬁc). Reverse transcription was performed using 1 µg of total RNA, oligo(dT) primer (Promega) and M-MLV reverse transcriptase (Promega). Real-time PCR was carried out using SybrGreen Master Mix (A&A Biotechnology) with the following cycling parameters: denaturation at 95º C for 20 sec, annealing at 62º C for 20 sec, and elongation at 72º C for 30 sec. Gene expression was normalized to elongation factor-2 (EF2) and the relative transcript levels were quantiﬁed using the 2^-ΔΔCt^ method. The following sequences of primers (Genomed) were used. *Ef2*, Forward: GACATCACCAAGGGTGTGCAG and Reverse: TTCAGCACACTGGCATAGAGGC; *Mcpip1*, Forward: CAGCCTCGACCAGATGTGCC and Reverse: CAGCCGCTCCTCGATGAAGC.

Western blotting

### Primary hepatocytes, Bone Marrow-derived Macrophages (BMDM), and tissue samples obtained from mice were lysed in RIPA buﬀer (25 mM Tris-HCl, pH 7.6; 150 mM NaCl; 1% sodium deoxycholate; 0.1% SDS) supplemented with Complete Protease Inhibitor Cocktail (Roche) and PhosSTOP Phosphatase Inhibitor Cocktail (Roche). Protein concentration was assessed using a bicinchoninic acid assay. Next, 20 μg of protein was separated on a 10% SDS-PAGE gel and transferred to a PVDF membrane (Millipore). The membrane was blocked for 1 h with 5% skim milk and incubated with primary antibodies overnight at 4º C. After incubation with secondary antibodies conjugated with HRP (1 h at room temperature), chemiluminescence was visualized using ECL^™^ Select Western Blotting Detection Reagent (GE Healthcare) in a MicroChemi chemiluminescence detector (BioRad). The following antibodies were used: rabbit anti-MCPIP1 (1:1000; Genetex), mouse anti-β-actin (1:4000; Sigma), peroxidase-conjugated anti-rabbit (1:30000; Cell Signaling) and peroxidase conjugated anti-mouse (1:20000; BD).

**Supplementary Figure legends**

**Fig. S1. Verification of *Zc3h12a* deletion in Mcpip1^AlbKO^ mice.**

(A) Schematic of LoxP site localization within the *Zc3h12a* gene; (B) Representative results of Mcpip1^fl/fl^ and Mcpip1^AlbKO^ mouse genotyping; (C) Expression of the *Zc3h12a* gene in various tissues; (D) Analysis of Mcpip1 protein level by western blotting; (E) Mass, (F) liver to body ratio, and (G) spleen to body ratio of 24-week-old Mcpip1^fl/fl^ (n=6) and Mcpip1^AlbKO^ (n=9) mice, *p < 0.05 vs. Mcpip1^fl/fl^.

**Fig. S2. Liver histology of Mcpip1^fl/fl^ mice.**

Representative hematoxylin & eosin staining (HE) and collagen staining (PSR, picrosirius red) of livers from 52-week-old male Mcpip1^fl/fl^ mice. Mcpip1^fl/fl^ mice of all ages had no pathological change in the liver. (A,C) liver parenchyma and (B,D) portal area. *, bile duct; V, vein; A, artery; CV, central vein. Magnification 200x, HE (A,D) and PSR (C,D) staining.

**Fig. S3. Progression of periportal pathology indicated via bile duct proliferation, inflammation, and fibrosis in Mcpip1^AlbKO^ male mice (6 to 52-week-old).**

Extensive bile duct hyperplasia with inflammatory infiltration and fibrosis were observed within the portal area in 6-week-old mice. These processes were temporarily decreased in the middle time points but were upregulated again in 52-week-old mice. Magnification 200x, HE (A-D) and PSR (E-H) staining (A, artery; V, vein; *, bile ducts). HE, hematoxylin and eosin; PSR, picrosirius red.

**Fig. S4. Mcpip1-dependent primary biliary cholangitis develops apart from systemic inflammation.**

Concentrations of selected cytokines and chemokines in plasma of (A) 6-week and (C) 24-week-old Mcpip1^fl/fl^, Mcpip1^LysMKO^, and Mcpip1^AlbKO^ mice; Total IgG and IgM in plasma of (B) 6-week and (D) 24-week-old mice presented as fold increase over Mcpip1^fl/fl^ controls; Production of (E) PGD_2_, (F) TXB_2_ and (G) 6-ketoPGF_1α_ by liver sinusoidal endothelial cells isolated form 24-week-old Mcpip1^fl/fl^ and knockout mice. ND, not detected. Data represent mean ± SEM, *p < 0.05 vs. Mcpip1 fl/fl.

**Fig. S5. Primary biliary cholangitis development is not intensified in female Mcpip1^AlbKO^ mice.**

(A) Amount of collagen in 6 and 24-week-old male (n=10) and female (n=10) Mcpip1^AlbKO^ mice, (B) Percentage of Mcpip1^AlbKO^ male and female mice with higher concentration of anti-PDC-E2 antimitochondrial autoantibodies and anti-gp-210 antinuclear autoantibodies compared to mean concentration in control Mcpip1^fl/fl^ counterparts.

**Fig. S6. Comparison of gene expression profiles of 6 and 24-week-old Mcpip1^AlbKO^ mice.**

GO enrichment analysis of genes differentially expressed in 6 and 24-week-old Mcpip1-depleted primary hepatocytes. Scale is the –log10(P-value) of the enrichment score (p-value < 0.05). (A) pathways induced and (B) inhibited in 6-week-old vs. 24-week-old Mcpip1^AlbKO^ hepatocytes; (C) Venn diagram presenting the number of transcripts significantly up- and downregulated (adj. P-value < 0.05) in 24-week-old Mcpip1^AlbKO^ samples (24w Mcpip1^AlbKO^ up/down) and 6-week-old Mcpip1^fl/fl^ (6w Mcpip1^fl/fl^ up/down) in comparison to 6-week-old Mcpip1^AlbKO^ samples.

**Figure S1.**

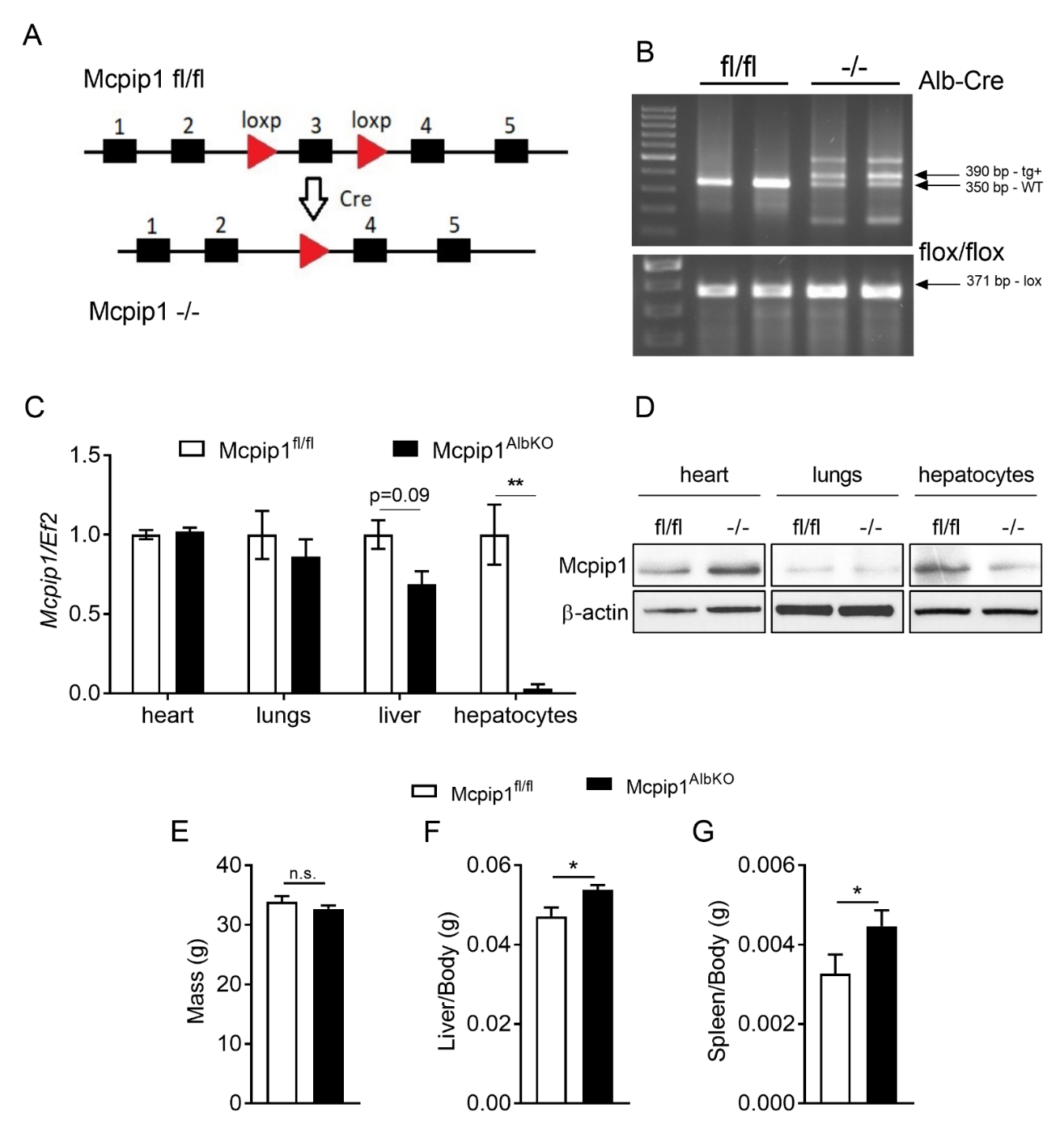

**Figure S2.**

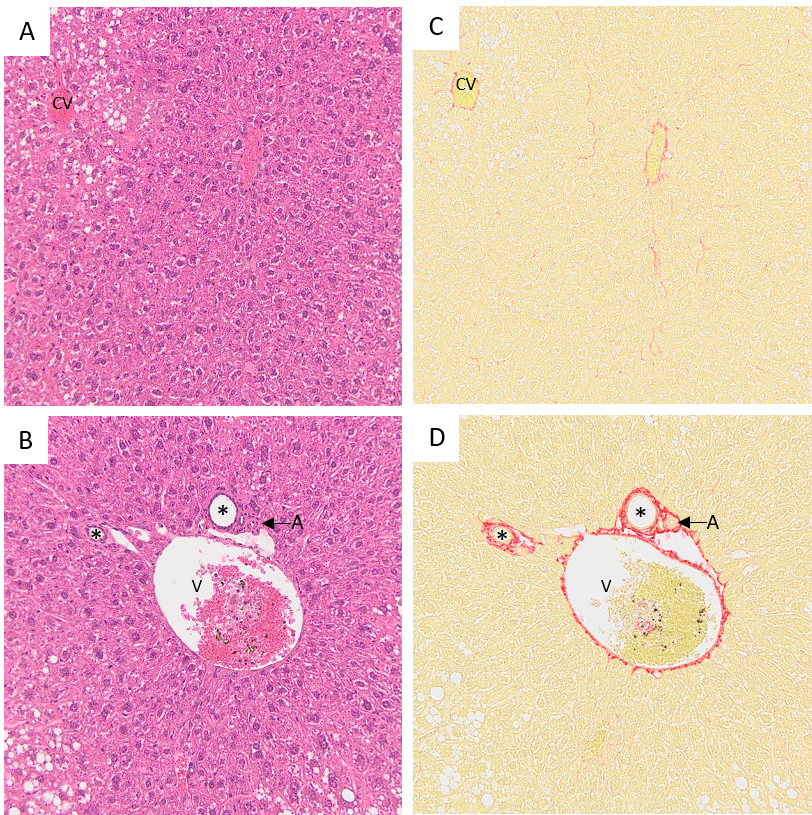

**Figure S3.**

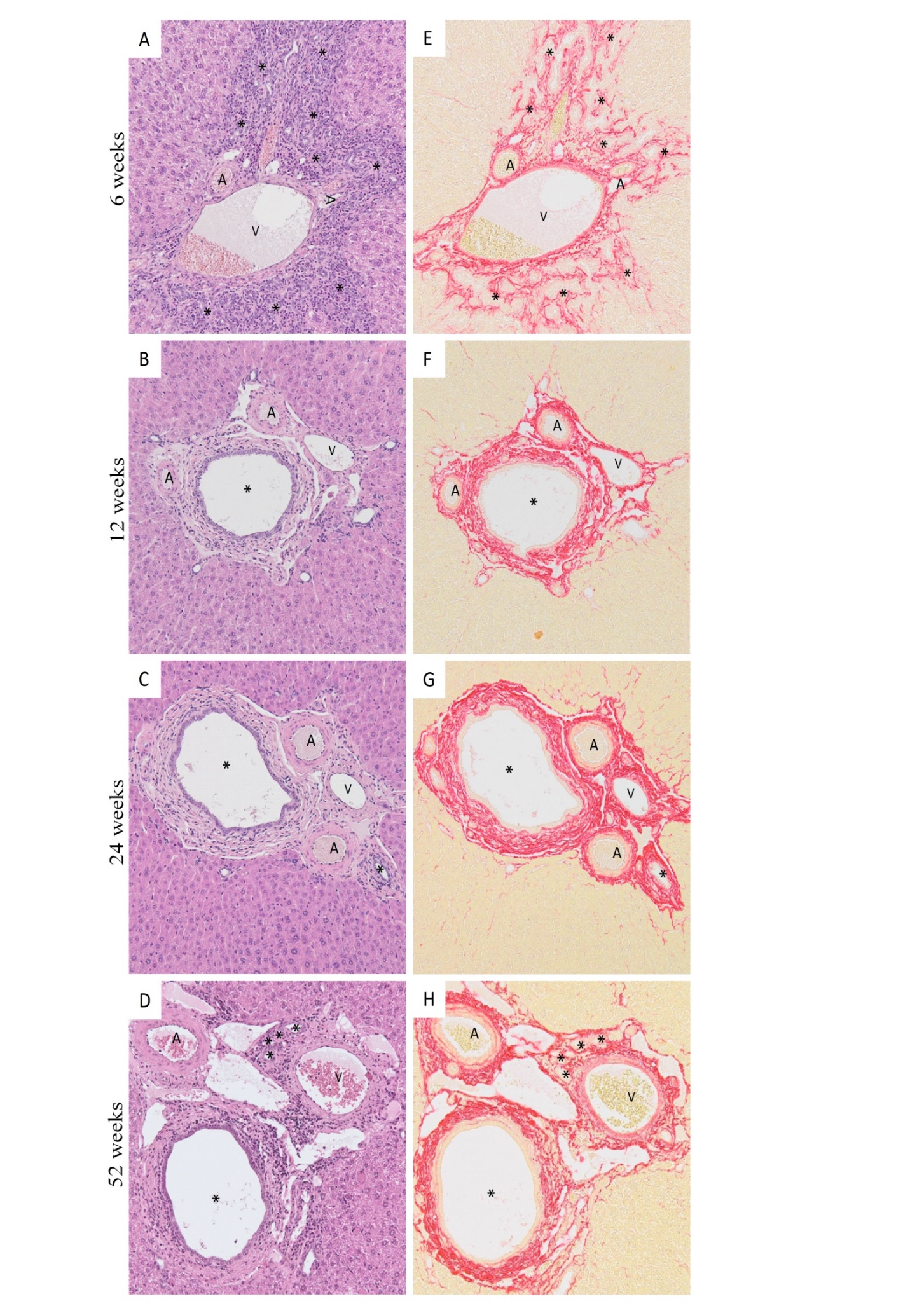

**Figure S4.**

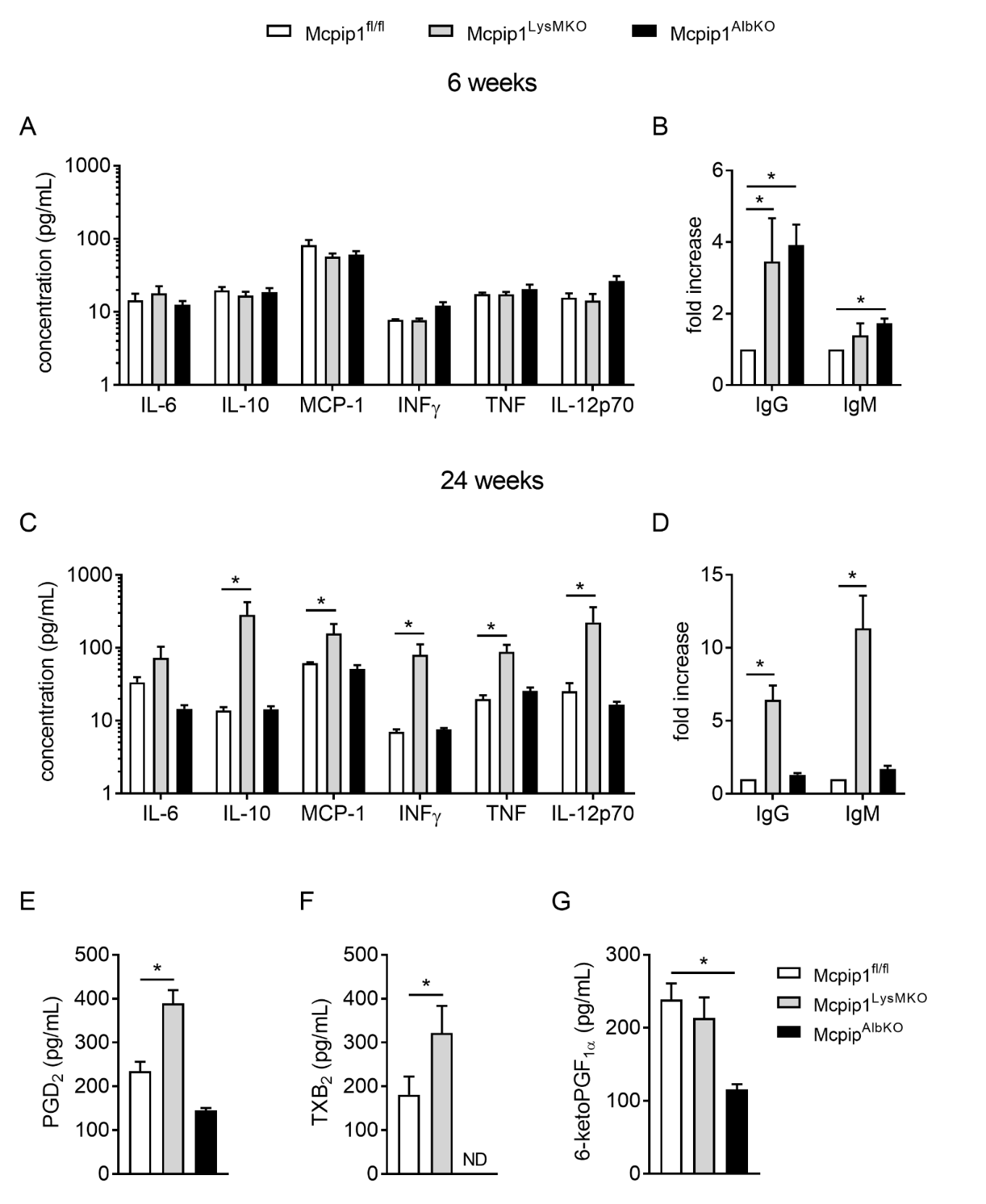

**Figure S5.**

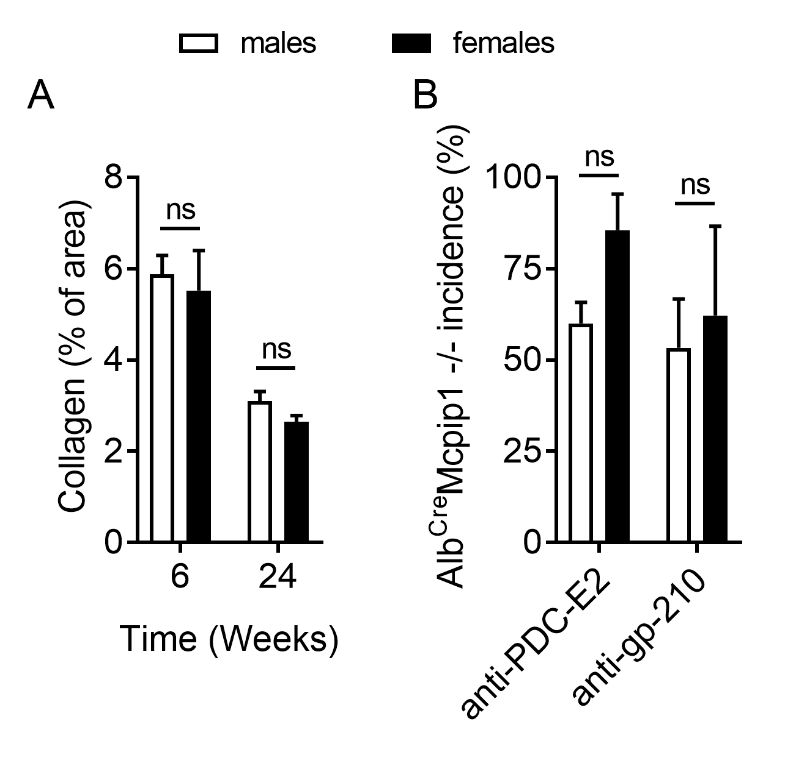

**Figure S6.**

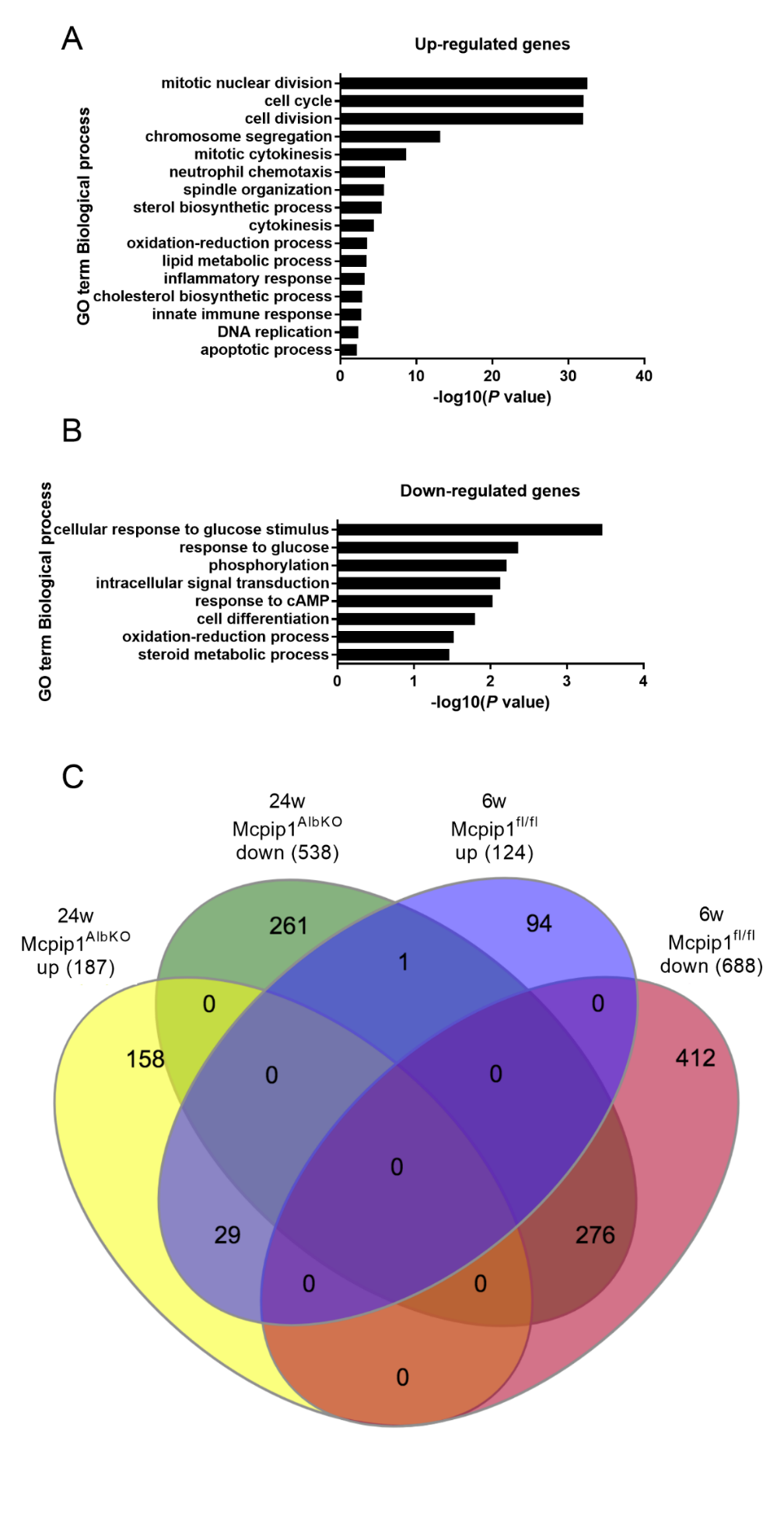

**Supplementary Tables**

**Table S1.** Genes differentially expressed in Mcpip1^fl/fl^ and Mcpip1^AlbKO^ hepatocytes collected from 6 weeks old mice.

| **Gene** | **Fold change** | **P value** |
| --- | --- | --- |
| *Ntrk2* | 33,46 | 8,308E-39 |
| *Mmp9* | 16,51 | 3,408E-22 |
| *Scd2* | 16,37 | 8,696E-29 |
| *Cxcl14* | 16,23 | 3,991E-20 |
| *Retnlg* | 15,78 | 9,565E-22 |
| *Cd14* | 15,32 | 8,749E-24 |
| *Cd44* | 11,55 | 1,370E-16 |
| *Mfge8* | 11,00 | 2,985E-19 |
| *Nid1* | 10,65 | 9,693E-25 |
| *Cybb* | 10,46 | 3,052E-16 |
| *Sirpa* | 10,38 | 9,043E-22 |
| *Lyz2* | 9,50 | 1,267E-25 |
| *Cd63* | 9,43 | 4,533E-25 |
| *Cd52* | 9,37 | 1,358E-16 |
| *Gabrb3* | 8,96 | 3,777E-12 |
| *Vim* | 8,88 | 9,012E-15 |
| *Ednrb* | 8,31 | 7,713E-12 |
| *Bicc1* | 8,18 | 1,227E-10 |
| *Ly6d* | 7,93 | 6,056E-10 |
| *Lcn2* | 7,89 | 1,485E-17 |
| *Tyrobp* | 7,84 | 4,950E-13 |
| *Cxcr4* | 7,46 | 7,896E-10 |
| *Cdkn3* | 7,26 | 2,880E-10 |
| *Gipc2* | 6,95 | 7,274E-09 |
| *H2-Aa* | 6,89 | 4,738E-12 |
| *Armcx4* | 6,80 | 1,101E-08 |
| *Slpi* | 6,65 | 1,648E-08 |
| *Cdk1* | 6,57 | 2,290E-15 |
| *Ccl6* | 6,52 | 1,282E-09 |
| *Msn* | 6,48 | 9,012E-15 |
| *Adam23* | 6,37 | 9,678E-13 |
| *Ncapg* | 6,35 | 1,465E-09 |
| *Cib3* | 6,18 | 3,988E-14 |
| *Cenpf* | 6,05 | 1,260E-11 |
| *Slc22a27* | 6,05 | 1,613E-07 |
| *Birc5* | 6,02 | 4,562E-10 |
| *Clec4d* | 6,00 | 2,458E-07 |
| *Prom1* | 5,97 | 2,417E-07 |
| *Alox5ap* | 5,88 | 2,469E-07 |
| *Ncf1* | 5,88 | 8,842E-08 |
| *Racgap1* | 5,81 | 1,865E-08 |
| *Kif20a* | 5,81 | 7,214E-20 |
| *Nuf2* | 5,80 | 8,918E-08 |
| *Coro1a* | 5,78 | 2,361E-07 |
| *Cyp2a22* | 5,78 | 3,299E-08 |
| *Golm1* | 5,68 | 4,734E-12 |
| *Cd74* | 5,66 | 9,987E-16 |
| *Pls1* | 5,66 | 1,777E-11 |
| *D17H6S56E-5* | 5,64 | 1,424E-10 |
| *Iqgap1* | 5,62 | 3,052E-16 |
| *Hdc* | 5,59 | 7,576E-07 |
| *Selplg* | 5,46 | 1,452E-06 |
| *Ccna2* | 5,45 | 7,046E-09 |
| *Sparc* | 5,39 | 1,511E-14 |
| *Rhoj* | 5,39 | 9,212E-08 |
| *Adrb2* | 5,37 | 2,775E-07 |
| *Pla2g7* | 5,35 | 1,489E-06 |
| *Sod3* | 5,35 | 4,131E-07 |
| *Serpinb6a* | 5,30 | 3,163E-12 |
| *Ifi27l2b* | 5,16 | 3,017E-11 |
| *Itgb2* | 5,14 | 1,465E-07 |
| *Ly6a* | 5,10 | 2,406E-09 |
| *Fam124a* | 5,09 | 7,586E-11 |
| *Stk10* | 5,08 | 6,209E-07 |
| *Snhg18* | 5,07 | 1,579E-07 |
| *Cd300lf* | 5,06 | 5,286E-06 |
| *Itgam* | 5,05 | 4,904E-06 |
| *H2-Q1* | 5,04 | 2,599E-09 |
| *Rgs2* | 5,04 | 4,087E-07 |
| *Tmsb4x* | 5,00 | 1,064E-10 |
| *Spink1* | 4,97 | 1,382E-06 |
| *Laptm5* | 4,96 | 7,898E-08 |
| *Rgs19* | 4,94 | 7,123E-06 |
| *Folr2* | 4,93 | 2,361E-07 |
| *Csf3r* | 4,88 | 9,012E-06 |
| *Fbn1* | 4,86 | 9,621E-06 |
| *Spp1* | 4,84 | 5,108E-06 |
| *Sdk1* | 4,82 | 9,684E-06 |
| *Lrtm2* | 4,81 | 1,135E-05 |
| *Fam83d* | 4,77 | 1,489E-06 |
| *Nckap1l* | 4,76 | 1,027E-05 |
| *Abcb1a* | 4,75 | 1,521E-06 |
| *Cd36* | 4,74 | 7,241E-10 |
| *Glipr2* | 4,74 | 1,301E-05 |
| *Fxyd5* | 4,74 | 4,015E-06 |
| *Tmeff2* | 4,73 | 5,738E-06 |
| *Lsp1* | 4,70 | 1,135E-05 |
| *Ppl* | 4,70 | 7,605E-07 |
| *Npdc1* | 4,66 | 4,452E-07 |
| *Slc26a10* | 4,66 | 2,305E-06 |
| *Pkm* | 4,63 | 1,101E-07 |
| *Lcp2* | 4,63 | 2,391E-06 |
| *Epdr1* | 4,63 | 1,918E-05 |
| *Entpd1* | 4,61 | 1,301E-05 |
| *Ptprc* | 4,61 | 1,752E-05 |
| *Slfn4* | 4,61 | 1,801E-05 |
| *Apbb1ip* | 4,58 | 1,310E-05 |
| *Ccdc80* | 4,55 | 6,345E-06 |
| *Anxa1* | 4,54 | 2,710E-05 |
| *Cyp4f16* | 4,52 | 7,065E-11 |
| *Ctla2b* | 4,51 | 9,105E-06 |
| *Prc1* | 4,48 | 4,904E-06 |
| *Aurka* | 4,47 | 4,356E-08 |
| *Srgn* | 4,47 | 1,942E-06 |
| *Cxcr2* | 4,46 | 3,592E-05 |
| *Slc13a5* | 4,40 | 2,083E-09 |
| *Abi2* | 4,37 | 7,164E-06 |
| *Cdc20* | 4,35 | 4,484E-07 |
| *Nfam1* | 4,34 | 2,330E-05 |
| *Cd5l* | 4,34 | 9,271E-08 |
| *Wfdc17* | 4,33 | 5,542E-06 |
| *Sybu* | 4,32 | 4,528E-05 |
| *H2-Eb1* | 4,31 | 1,801E-05 |
| *Arhgap30* | 4,30 | 6,075E-05 |
| *Ngfrap1* | 4,28 | 3,734E-05 |
| *Kif22* | 4,27 | 3,742E-05 |
| *S100a9* | 4,25 | 6,802E-05 |
| *Fmo2* | 4,24 | 1,789E-08 |
| *Kif23* | 4,23 | 7,181E-05 |
| *Prelp* | 4,19 | 7,094E-07 |
| *Sorl1* | 4,19 | 5,975E-05 |
| *Rab27a* | 4,19 | 1,524E-06 |
| *Cmtm3* | 4,18 | 1,238E-05 |
| *Kif2c* | 4,17 | 5,596E-05 |
| *Mki67* | 4,17 | 9,386E-07 |
| *Alpk1* | 4,14 | 1,092E-04 |
| *Gcnt1* | 4,13 | 1,092E-04 |
| *Csf2rb2* | 4,12 | 1,099E-04 |
| *Adgre5* | 4,12 | 7,164E-06 |
| *Fcgr4* | 4,11 | 1,100E-04 |
| *Epsti1* | 4,09 | 1,214E-04 |
| *Cd93* | 4,08 | 1,259E-05 |
| *Mcam* | 4,07 | 9,677E-06 |
| *Kit* | 4,07 | 1,230E-04 |
| *1700112E06Rik* | 4,07 | 7,445E-05 |
| *Osmr* | 4,05 | 7,804E-05 |
| *Rcsd1* | 4,05 | 8,149E-05 |
| *Tuba8* | 4,04 | 8,572E-07 |
| *Pfkp* | 4,03 | 1,492E-04 |
| *Arhgdib* | 4,03 | 3,184E-06 |
| *Sdr16c5* | 4,02 | 1,301E-04 |
| *Arhgap11a* | 4,02 | 6,807E-06 |
| *Rac2* | 4,00 | 1,671E-04 |
| *Plvap* | 3,99 | 3,114E-05 |
| *Hmmr* | 3,98 | 3,298E-05 |
| *Trim59* | 3,95 | 1,727E-04 |
| *Dcn* | 3,95 | 1,942E-06 |
| *Hspg2* | 3,94 | 2,736E-09 |
| *Csrp1* | 3,94 | 5,421E-07 |
| *Nusap1* | 3,93 | 9,506E-06 |
| *Prtn3* | 3,91 | 2,105E-04 |
| *Cep55* | 3,91 | 1,039E-04 |
| *Vsig4* | 3,90 | 6,333E-05 |
| *Slfn1* | 3,89 | 2,209E-04 |
| *Serpina3g* | 3,87 | 7,700E-06 |
| *Cdca5* | 3,86 | 1,719E-05 |
| *Tmem71* | 3,85 | 7,014E-05 |
| *Jam2* | 3,85 | 5,236E-07 |
| *Tacc3* | 3,84 | 2,943E-05 |
| *Ets1* | 3,81 | 1,371E-05 |
| *Gucy1a3* | 3,81 | 3,442E-04 |
| *Cd300a* | 3,80 | 2,664E-04 |
| *Blnk* | 3,79 | 6,543E-06 |
| *Dpysl2* | 3,78 | 6,327E-06 |
| *Smpd3* | 3,76 | 3,486E-04 |
| *Kif14* | 3,74 | 3,571E-04 |
| *Cav2* | 3,73 | 1,789E-08 |
| *Nlrp3* | 3,73 | 3,901E-04 |
| *Dpysl3* | 3,72 | 2,805E-04 |
| *Nedd9* | 3,71 | 6,978E-07 |
| *Pilra* | 3,69 | 4,096E-04 |
| *Gpsm3* | 3,68 | 1,327E-04 |
| *Tgfbi* | 3,68 | 1,141E-04 |
| *Fcgr3* | 3,66 | 2,679E-04 |
| *Rab3b* | 3,66 | 4,980E-04 |
| *Ncf2* | 3,66 | 5,813E-04 |
| *Kif18b* | 3,65 | 3,236E-04 |
| *Ccdc120* | 3,63 | 7,399E-05 |
| *Csf1r* | 3,63 | 4,791E-04 |
| *Abcc5* | 3,62 | 6,374E-04 |
| *Slc11a1* | 3,62 | 7,504E-05 |
| *Lrrc25* | 3,62 | 6,589E-04 |
| *Gsn* | 3,62 | 6,609E-04 |
| *Emp3* | 3,61 | 5,893E-04 |
| *Cdc42ep5* | 3,61 | 7,504E-05 |
| *Calcrl* | 3,60 | 7,381E-06 |
| *C1qtnf1* | 3,57 | 7,123E-06 |
| *Akr1b8* | 3,56 | 3,657E-04 |
| *Tnfrsf11b* | 3,56 | 7,949E-04 |
| *Col1a2* | 3,56 | 7,129E-04 |
| *Ccnb2* | 3,56 | 5,820E-05 |
| *Il1a* | 3,55 | 5,967E-04 |
| *Dab2* | 3,53 | 4,179E-06 |
| *Fmnl1* | 3,52 | 8,973E-04 |
| *Atp1a3* | 3,51 | 7,609E-04 |
| *Parvb* | 3,51 | 9,614E-04 |
| *Fgl2* | 3,50 | 9,614E-04 |
| *Tmem173* | 3,49 | 8,020E-04 |
| *Clec4g* | 3,49 | 5,286E-06 |
| *Gpihbp1* | 3,48 | 6,343E-05 |
| *Top2a* | 3,48 | 3,895E-04 |
| *Efemp2* | 3,47 | 1,055E-03 |
| *Bub1b* | 3,47 | 4,231E-04 |
| *Adgre4* | 3,47 | 7,340E-04 |
| *Elmo1* | 3,46 | 5,036E-04 |
| *Adgra2* | 3,46 | 9,198E-04 |
| *Ckap2l* | 3,46 | 1,533E-04 |
| *Nek2* | 3,45 | 4,096E-07 |
| *Thbd* | 3,44 | 1,005E-03 |
| *Spdl1* | 3,44 | 7,602E-04 |
| *Selp* | 3,43 | 1,020E-03 |
| *Armcx2* | 3,43 | 1,180E-03 |
| *Cdh5* | 3,42 | 1,489E-06 |
| *Klf2* | 3,42 | 3,734E-05 |
| *Ppp1r9b* | 3,42 | 7,146E-04 |
| *Ctss* | 3,42 | 6,390E-04 |
| *Ifi205* | 3,41 | 1,355E-03 |
| *Rasgrp3* | 3,41 | 4,530E-04 |
| *Slc9a9* | 3,40 | 5,525E-04 |
| *Cntln* | 3,40 | 1,055E-03 |
| *Orm2* | 3,40 | 4,015E-06 |
| *Pglyrp1* | 3,39 | 1,466E-03 |
| *Aurkb* | 3,38 | 6,106E-04 |
| *Fcna* | 3,37 | 2,942E-04 |
| *Espl1* | 3,37 | 1,520E-03 |
| *Pak6* | 3,37 | 1,180E-03 |
| *Melk* | 3,36 | 1,520E-03 |
| *Col14a1* | 3,36 | 8,344E-04 |
| *Plk1* | 3,36 | 5,855E-04 |
| *Chst15* | 3,34 | 5,763E-05 |
| *Plekho2* | 3,34 | 7,949E-04 |
| *Col5a2* | 3,34 | 1,754E-03 |
| *Kifc1* | 3,33 | 7,752E-04 |
| *Trim47* | 3,33 | 1,355E-03 |
| *Prrg4* | 3,33 | 9,070E-04 |
| *Nrm* | 3,32 | 8,665E-04 |
| *Fcer1g* | 3,32 | 7,708E-04 |
| *Marco* | 3,32 | 1,827E-03 |
| *Gngt2* | 3,32 | 6,106E-04 |
| *Ift57* | 3,31 | 1,214E-04 |
| *Slc25a4* | 3,30 | 1,719E-05 |
| *Pygb* | 3,30 | 8,797E-04 |
| *Rragd* | 3,30 | 1,529E-03 |
| *Ect2* | 3,29 | 1,238E-05 |
| *Tspan7* | 3,29 | 8,361E-06 |
| *Plekhg1* | 3,29 | 3,980E-04 |
| *Anxa3* | 3,28 | 1,779E-03 |
| *Fam111a* | 3,28 | 1,885E-03 |
| *Hebp2* | 3,28 | 1,774E-03 |
| *H2-Ab1* | 3,27 | 1,349E-03 |
| *Spi1* | 3,27 | 1,827E-03 |
| *Cxcl2* | 3,27 | 1,820E-03 |
| *Gm33543* | 3,26 | 1,194E-07 |
| *Itpkb* | 3,26 | 3,571E-04 |
| *Il1rn* | 3,26 | 7,340E-04 |
| *Flt4* | 3,25 | 2,779E-05 |
| *Dst* | 3,25 | 5,852E-07 |
| *Knstrn* | 3,25 | 5,293E-04 |
| *Clec7a* | 3,24 | 8,415E-04 |
| *Bgn* | 3,24 | 1,857E-04 |
| *Cyb561* | 3,24 | 5,542E-06 |
| *Mcemp1* | 3,23 | 1,859E-03 |
| *Slc44a2* | 3,23 | 1,124E-03 |
| *Mtmr11* | 3,22 | 2,654E-03 |
| *Lhfp* | 3,22 | 1,903E-03 |
| *Osbpl3* | 3,22 | 2,664E-03 |
| *Slc15a3* | 3,22 | 2,510E-03 |
| *Ctgf* | 3,21 | 2,759E-03 |
| *Tfec* | 3,21 | 2,111E-03 |
| *Kif20b* | 3,20 | 4,644E-04 |
| *Ntf3* | 3,20 | 2,232E-03 |
| *Rhbdf1* | 3,19 | 1,861E-04 |
| *Plaur* | 3,19 | 2,760E-03 |
| *Serpinb8* | 3,19 | 1,099E-04 |
| *Serpina7* | 3,19 | 8,146E-05 |
| *Veph1* | 3,18 | 2,564E-03 |
| *Ccl2* | 3,18 | 3,078E-03 |
| *Psrc1* | 3,17 | 3,075E-03 |
| *Ddr1* | 3,17 | 3,121E-03 |
| *Evl* | 3,17 | 3,192E-03 |
| *Iqgap3* | 3,17 | 1,779E-03 |
| *Sh3bgrl3* | 3,16 | 3,895E-04 |
| *Igf1r* | 3,16 | 2,988E-03 |
| *Parpbp* | 3,15 | 3,346E-03 |
| *Cd24a* | 3,15 | 3,389E-03 |
| *Adgrl4* | 3,15 | 1,328E-04 |
| *Ankle1* | 3,14 | 3,121E-03 |
| *Rftn1* | 3,13 | 3,653E-03 |
| *Cdca8* | 3,12 | 1,082E-03 |
| *Adgrg1* | 3,12 | 7,704E-04 |
| *Il16* | 3,12 | 2,732E-03 |
| *Arhgap25* | 3,12 | 3,790E-03 |
| *Cand2* | 3,12 | 1,975E-03 |
| *Coro2b* | 3,11 | 3,564E-03 |
| *Ccl3* | 3,11 | 3,239E-03 |
| *Siglece* | 3,10 | 3,957E-03 |
| *Itgax* | 3,09 | 3,567E-03 |
| *Tifa* | 3,09 | 2,935E-08 |
| *Abhd2* | 3,08 | 9,987E-16 |
| *Neurl1b* | 3,08 | 1,793E-03 |
| *Serpinh1* | 3,08 | 1,686E-03 |
| *Lysmd2* | 3,08 | 2,344E-03 |
| *Pde2a* | 3,08 | 4,428E-04 |
| *Cxcl16* | 3,08 | 2,673E-05 |
| *Wls* | 3,07 | 2,019E-03 |
| *Clec12a* | 3,07 | 4,553E-03 |
| *Gpx3* | 3,06 | 3,313E-03 |
| *Atp6v0d2* | 3,06 | 8,292E-04 |
| *5330417C22Rik* | 3,06 | 4,777E-03 |
| *Sytl5* | 3,06 | 2,558E-03 |
| *Shank3* | 3,05 | 4,777E-03 |
| *Flna* | 3,05 | 1,549E-03 |
| *Slc16a5* | 3,04 | 4,003E-04 |
| *Oasl2* | 3,04 | 2,325E-04 |
| *Nrg1* | 3,04 | 5,015E-03 |
| *Tpx2* | 3,04 | 2,439E-03 |
| *Plekha2* | 3,03 | 5,015E-03 |
| *Hao2* | 3,03 | 3,472E-03 |
| *Arhgef15* | 3,02 | 5,127E-03 |
| *Clec4f* | 3,01 | 9,167E-04 |
| *Slc41a3* | 3,01 | 3,484E-03 |
| *Fstl1* | 3,00 | 5,561E-03 |
| *Plscr4* | 3,00 | 5,880E-03 |
| *Vav1* | 2,99 | 6,028E-03 |
| *Ptprb* | 2,99 | 1,492E-04 |
| *Afap1l1* | 2,99 | 5,525E-03 |
| *Plek* | 2,99 | 6,030E-03 |
| *Plxnc1* | 2,98 | 6,978E-04 |
| *Per1* | 2,98 | 3,386E-05 |
| *Uhrf1* | 2,98 | 4,574E-03 |
| *Adgre1* | 2,97 | 6,270E-03 |
| *Cytip* | 2,97 | 5,561E-03 |
| *Ube2c* | 2,97 | 5,922E-03 |
| *Cnn2* | 2,97 | 3,084E-03 |
| *Pde3a* | 2,97 | 5,493E-03 |
| *AI607873* | 2,96 | 4,112E-03 |
| *Uap1l1* | 2,96 | 1,414E-04 |
| *Aqp1* | 2,96 | 1,224E-04 |
| *Sdc3* | 2,96 | 1,154E-03 |
| *Gja1* | 2,95 | 1,582E-03 |
| *Sirpb1b* | 2,95 | 5,004E-03 |
| *Sp100* | 2,95 | 1,055E-03 |
| *Igfbp7* | 2,95 | 4,021E-04 |
| *Tinag* | 2,95 | 6,374E-03 |
| *Stxbp1* | 2,95 | 7,089E-03 |
| *Slc38a4* | 2,94 | 1,327E-04 |
| *Cyfip2* | 2,94 | 6,493E-03 |
| *Plxnd1* | 2,94 | 2,323E-03 |
| *Oas3* | 2,94 | 6,561E-03 |
| *Kif11* | 2,93 | 3,357E-03 |
| *Ptgs1* | 2,93 | 5,422E-03 |
| *Shc2* | 2,93 | 5,255E-03 |
| *Pbk* | 2,93 | 5,593E-03 |
| *Mastl* | 2,92 | 3,393E-03 |
| *Vcam1* | 2,92 | 5,043E-03 |
| *S100a8* | 2,92 | 5,525E-03 |
| *Bmp2* | 2,92 | 1,479E-03 |
| *Clec4n* | 2,92 | 7,028E-03 |
| *Cd37* | 2,92 | 8,075E-03 |
| *Kalrn* | 2,91 | 5,270E-03 |
| *Emp2* | 2,90 | 5,801E-05 |
| *Amotl1* | 2,90 | 2,664E-03 |
| *Nt5c3b* | 2,90 | 4,208E-05 |
| *Neil3* | 2,90 | 8,324E-03 |
| *Ltb* | 2,89 | 8,075E-03 |
| *Cd38* | 2,89 | 1,744E-03 |
| *Ntn4* | 2,88 | 1,496E-03 |
| *Casp1* | 2,88 | 8,299E-03 |
| *Slc43a2* | 2,88 | 2,289E-03 |
| *Fam198b* | 2,88 | 7,433E-03 |
| *Fabp4* | 2,88 | 3,084E-03 |
| *Shcbp1* | 2,88 | 2,492E-03 |
| *Calml4* | 2,88 | 6,211E-03 |
| *Ecm1* | 2,87 | 1,200E-03 |
| *Eda2r* | 2,86 | 8,075E-03 |
| *Samsn1* | 2,86 | 8,550E-03 |
| *Fyn* | 2,85 | 4,627E-03 |
| *Tlr4* | 2,85 | 9,313E-03 |
| *Gdf2* | 2,84 | 9,908E-03 |
| *Dsn1* | 2,84 | 6,109E-03 |
| *Cdca2* | 2,84 | 6,493E-03 |
| *Syce2* | 2,83 | 8,052E-03 |
| *Trem1* | 2,83 | 8,712E-03 |
| *Dock2* | 2,83 | 1,024E-02 |
| *Vwf* | 2,82 | 8,826E-03 |
| *Csf2ra* | 2,81 | 3,544E-03 |
| *Morc4* | 2,80 | 5,922E-03 |
| *Defb1* | 2,80 | 3,943E-03 |
| *Dmpk* | 2,80 | 7,135E-03 |
| *C3ar1* | 2,80 | 1,087E-02 |
| *Cd101* | 2,79 | 9,969E-03 |
| *Fmo3* | 2,79 | 7,089E-03 |
| *Gm33447* | 2,79 | 9,847E-03 |
| *Esam* | 2,79 | 2,611E-03 |
| *Il1f9* | 2,79 | 1,123E-02 |
| *Syngr4* | 2,78 | 1,037E-02 |
| *Tfpi* | 2,78 | 1,790E-04 |
| *Elf4* | 2,77 | 1,150E-02 |
| *Ncapd2* | 2,77 | 7,704E-04 |
| *Eng* | 2,77 | 3,571E-04 |
| *Il1b* | 2,77 | 1,024E-02 |
| *Itga4* | 2,77 | 1,349E-02 |
| *Col3a1* | 2,76 | 1,446E-02 |
| *Gpr182* | 2,76 | 1,862E-03 |
| *Casc5* | 2,75 | 1,122E-02 |
| *Ly6e* | 2,75 | 1,190E-06 |
| *Tgfb1* | 2,75 | 1,279E-02 |
| *Rassf2* | 2,74 | 1,146E-02 |
| *Elovl7* | 2,74 | 1,019E-02 |
| *Myb* | 2,74 | 9,969E-03 |
| *Kdr* | 2,73 | 1,466E-03 |
| *Atp11a* | 2,73 | 1,140E-03 |
| *Mndal* | 2,73 | 8,781E-03 |
| *Colec11* | 2,73 | 1,779E-03 |
| *Armcx1* | 2,73 | 1,376E-02 |
| *Eif4e3* | 2,73 | 6,030E-03 |
| *Ngf* | 2,73 | 1,614E-02 |
| *Cygb* | 2,73 | 1,507E-02 |
| *Col4a2* | 2,72 | 2,284E-04 |
| *Bmper* | 2,72 | 1,455E-02 |
| *Ifi203* | 2,72 | 9,583E-03 |
| *Ccl24* | 2,72 | 1,661E-02 |
| *Wee1* | 2,72 | 8,397E-05 |
| *Tlr8* | 2,72 | 1,629E-02 |
| *Irak3* | 2,70 | 1,538E-02 |
| *Axl* | 2,70 | 1,759E-02 |
| *Oit3* | 2,69 | 1,155E-05 |
| *Adgrf5* | 2,69 | 1,873E-03 |
| *Spats2l* | 2,69 | 1,864E-02 |
| *Tmcc2* | 2,68 | 1,901E-02 |
| *Fgd1* | 2,68 | 6,561E-03 |
| *Rcn1* | 2,68 | 1,507E-02 |
| *Rbp1* | 2,68 | 2,163E-07 |
| *Mrc1* | 2,68 | 1,997E-03 |
| *Tnfrsf10b* | 2,67 | 9,331E-03 |
| *Ms4a6c* | 2,66 | 1,583E-02 |
| *Diaph3* | 2,66 | 1,499E-02 |
| *B430306N03Rik* | 2,66 | 1,668E-02 |
| *Vsig10* | 2,65 | 1,805E-02 |
| *Sh3kbp1* | 2,65 | 2,104E-02 |
| *Ank3* | 2,65 | 1,453E-04 |
| *Tspan15* | 2,65 | 1,906E-02 |
| *Mxra8* | 2,64 | 2,116E-02 |
| *Gpx7* | 2,64 | 1,496E-03 |
| *Stab1* | 2,64 | 3,881E-03 |
| *Fads3* | 2,63 | 1,583E-02 |
| *Col4a1* | 2,63 | 4,080E-04 |
| *Robo4* | 2,63 | 1,351E-02 |
| *Emilin1* | 2,63 | 1,049E-02 |
| *Acot9* | 2,62 | 2,574E-05 |
| *Wdfy4* | 2,62 | 2,314E-02 |
| *Fabp7* | 2,62 | 2,428E-02 |
| *Gdf10* | 2,62 | 2,201E-02 |
| *Bmp5* | 2,62 | 1,924E-02 |
| *Mlkl* | 2,62 | 1,479E-03 |
| *Cmklr1* | 2,61 | 2,341E-02 |
| *Klhl6* | 2,61 | 2,346E-02 |
| *Cdkn2c* | 2,61 | 1,752E-05 |
| *Hist1h3d* | 2,61 | 1,356E-02 |
| *Gbp2* | 2,61 | 2,494E-02 |
| *Lfng* | 2,60 | 2,543E-02 |
| *Ppp1r14a* | 2,60 | 3,550E-03 |
| *Cfp* | 2,60 | 1,629E-02 |
| *Mthfd1l* | 2,59 | 3,056E-03 |
| *Steap2* | 2,59 | 4,343E-04 |
| *Cdca3* | 2,58 | 1,949E-03 |
| *Ifi30* | 2,58 | 2,204E-04 |
| *Dok3* | 2,58 | 2,647E-02 |
| *Stab2* | 2,58 | 4,789E-04 |
| *Gal3st1* | 2,58 | 4,406E-03 |
| *Pdgfrb* | 2,57 | 2,821E-02 |
| *Rad54l* | 2,57 | 2,821E-02 |
| *Sept4* | 2,57 | 2,553E-02 |
| *Itih5* | 2,57 | 2,165E-04 |
| *Dchs1* | 2,56 | 2,927E-02 |
| *Gucy1b3* | 2,56 | 2,964E-02 |
| *Vopp1* | 2,55 | 2,584E-02 |
| *Sgol2a* | 2,55 | 2,151E-02 |
| *Vav3* | 2,55 | 2,317E-02 |
| *Ramp2* | 2,55 | 2,723E-02 |
| *Pde10a* | 2,55 | 2,333E-02 |
| *F8* | 2,54 | 1,610E-03 |
| *Ppp1r16b* | 2,54 | 3,099E-02 |
| *Ckap2* | 2,54 | 1,390E-02 |
| *Abcg1* | 2,54 | 1,385E-02 |
| *Lpcat2* | 2,54 | 3,009E-02 |
| *Ppp1r18* | 2,54 | 3,009E-02 |
| *Ppp1r3d* | 2,53 | 2,027E-02 |
| *Rgs10* | 2,53 | 2,924E-02 |
| *Dhrs9* | 2,53 | 7,300E-04 |
| *Ptgr1* | 2,53 | 9,561E-05 |
| *Clec1b* | 2,53 | 2,502E-02 |
| *Soga1* | 2,52 | 2,073E-02 |
| *Espn* | 2,52 | 3,349E-02 |
| *Arhgap31* | 2,52 | 2,465E-02 |
| *Prr11* | 2,52 | 2,520E-02 |
| *Pak1* | 2,52 | 2,873E-02 |
| *Col1a1* | 2,52 | 3,142E-02 |
| *Exoc3l4* | 2,52 | 3,142E-02 |
| *Rab31* | 2,52 | 3,009E-02 |
| *Gm11454* | 2,51 | 3,311E-02 |
| *Lair1* | 2,51 | 3,472E-02 |
| *Aldh3b1* | 2,51 | 2,082E-02 |
| *2010003K11Rik* | 2,51 | 1,108E-03 |
| *Entpd2* | 2,50 | 6,775E-04 |
| *Prex2* | 2,50 | 2,064E-02 |
| *Cerk* | 2,50 | 3,225E-02 |
| *Zc2hc1a* | 2,50 | 2,850E-02 |
| *Cd72* | 2,49 | 3,496E-02 |
| *Arhgef39* | 2,49 | 3,613E-02 |
| *Rab8b* | 2,49 | 3,520E-02 |
| *Timp3* | 2,49 | 1,012E-02 |
| *Smim24* | 2,49 | 3,579E-02 |
| *Tubb6* | 2,49 | 1,295E-04 |
| *Sell* | 2,49 | 3,248E-02 |
| *Fam49a* | 2,49 | 3,756E-02 |
| *Ackr3* | 2,48 | 3,451E-02 |
| *Pld4* | 2,48 | 3,286E-02 |
| *Slc5a1* | 2,47 | 3,518E-02 |
| *Msantd3* | 2,47 | 3,892E-02 |
| *Plpp1* | 2,47 | 2,089E-04 |
| *Adamts5* | 2,47 | 2,817E-02 |
| *Ccnd1* | 2,47 | 8,066E-05 |
| *Cln6* | 2,46 | 1,364E-02 |
| *Aldh18a1* | 2,46 | 4,023E-02 |
| *Anln* | 2,46 | 3,311E-02 |
| *Kntc1* | 2,45 | 4,183E-02 |
| *Prr15l* | 2,45 | 3,399E-02 |
| *Lrrc8c* | 2,45 | 2,554E-02 |
| *Arhgef6* | 2,45 | 3,951E-02 |
| *Gimap6* | 2,45 | 1,497E-03 |
| *Hgf* | 2,45 | 3,902E-02 |
| *Fam129a* | 2,45 | 3,908E-02 |
| *Map4k4* | 2,44 | 7,154E-03 |
| *Tff3* | 2,44 | 8,075E-03 |
| *Crim1* | 2,44 | 1,264E-04 |
| *AA986860* | 2,44 | 4,479E-02 |
| *Gucy1a2* | 2,44 | 4,064E-02 |
| *Hcst* | 2,43 | 3,248E-02 |
| *Rbms3* | 2,43 | 3,703E-02 |
| *Arhgap44* | 2,43 | 3,311E-02 |
| *Wnt2* | 2,43 | 4,611E-02 |
| *Slc25a24* | 2,43 | 3,504E-02 |
| *Tbx2* | 2,42 | 4,399E-02 |
| *Ccnf* | 2,42 | 6,898E-03 |
| *Rad51* | 2,42 | 3,137E-02 |
| *Fam102b* | 2,42 | 4,621E-02 |
| *Ttk* | 2,42 | 4,183E-02 |
| *Cd53* | 2,41 | 4,743E-02 |
| *Evc* | 2,41 | 4,810E-02 |
| *Slc37a1* | 2,41 | 1,590E-02 |
| *B4galt5* | 2,41 | 4,750E-03 |
| *Hk1* | 2,41 | 4,542E-02 |
| *Nipal1* | 2,40 | 3,703E-02 |
| *Dyrk3* | 2,40 | 2,989E-02 |
| *Pam* | 2,40 | 3,951E-02 |
| *Aspm* | 2,40 | 3,518E-02 |
| *Cd9* | 2,39 | 4,081E-09 |
| *Tagln2* | 2,39 | 3,128E-02 |
| *Zeb2* | 2,39 | 3,009E-02 |
| *Rgs4* | 2,39 | 4,187E-02 |
| *Themis2* | 2,39 | 4,480E-02 |
| *E330011O21Rik* | 2,39 | 3,395E-02 |
| *Arhgef16* | 2,39 | 2,211E-02 |
| *Abcd2* | 2,39 | 2,880E-03 |
| *Rtn4* | 2,39 | 5,454E-06 |
| *Chmp4c* | 2,39 | 3,579E-02 |
| *Tmem204* | 2,39 | 1,995E-02 |
| *Mylip* | 2,39 | 2,243E-02 |
| *Gprc5b* | 2,38 | 4,395E-02 |
| *Neurl3* | 2,38 | 4,810E-02 |
| *Gimap4* | 2,38 | 2,599E-02 |
| *Elk3* | 2,38 | 3,142E-02 |
| *Cers6* | 2,38 | 4,916E-04 |
| *Rrm2* | 2,37 | 3,084E-03 |
| *Plet1* | 2,37 | 3,305E-02 |
| *Mmgt2* | 2,37 | 2,817E-02 |
| *Colec10* | 2,37 | 4,640E-02 |
| *2810025M15Rik* | 2,36 | 1,867E-02 |
| *Tfrc* | 2,36 | 2,196E-03 |
| *Nfkbid* | 2,36 | 4,593E-02 |
| *Incenp* | 2,36 | 2,896E-02 |
| *Tcf24* | 2,36 | 2,964E-02 |
| *Cyp2a4* | 2,35 | 4,620E-02 |
| *Dnase1l3* | 2,35 | 3,484E-03 |
| *Ccr2* | 2,35 | 4,954E-02 |
| *Igfbp3* | 2,34 | 4,810E-02 |
| *Casp12* | 2,34 | 4,976E-02 |
| *Fli1* | 2,33 | 4,426E-02 |
| *AU021092* | 2,33 | 3,879E-02 |
| *Tinagl1* | 2,33 | 2,821E-02 |
| *Dnah5* | 2,33 | 3,804E-02 |
| *Tceal1* | 2,32 | 3,103E-02 |
| *B3gnt9* | 2,32 | 4,021E-02 |
| *Mad2l1* | 2,32 | 3,313E-03 |
| *Erg* | 2,31 | 4,954E-02 |
| *Ugt2b37* | 2,31 | 3,308E-03 |
| *Mfsd7c* | 2,31 | 3,313E-03 |
| *C1qb* | 2,31 | 4,935E-02 |
| *Gas6* | 2,30 | 1,224E-04 |
| *Tek* | 2,30 | 2,026E-02 |
| *Arvcf* | 2,30 | 4,889E-02 |
| *Fam198a* | 2,29 | 2,333E-02 |
| *Mvb12b* | 2,29 | 3,834E-02 |
| *Cotl1* | 2,29 | 1,981E-02 |
| *Arhgap19* | 2,29 | 2,494E-02 |
| *Tmem237* | 2,29 | 4,534E-04 |
| *Hexb* | 2,29 | 3,758E-05 |
| *Acod1* | 2,28 | 4,440E-02 |
| *Ccdc34* | 2,27 | 2,631E-02 |
| *Anxa5* | 2,26 | 1,292E-08 |
| *Cd300lg* | 2,25 | 4,362E-02 |
| *Pafah1b3* | 2,25 | 8,687E-03 |
| *Irf8* | 2,25 | 2,554E-02 |
| *Gls* | 2,25 | 3,552E-02 |
| *Ncam2* | 2,24 | 2,461E-02 |
| *Cep192* | 2,22 | 2,477E-02 |
| *Smc4* | 2,22 | 2,419E-04 |
| *App* | 2,21 | 1,834E-04 |
| *Rgl1* | 2,21 | 2,024E-02 |
| *Haus8* | 2,20 | 2,317E-02 |
| *Tmem43* | 2,20 | 2,855E-03 |
| *Epb41l2* | 2,19 | 2,893E-02 |
| *Plscr1* | 2,18 | 1,956E-04 |
| *Slc20a1* | 2,17 | 1,453E-04 |
| *Crtap* | 2,17 | 6,907E-03 |
| *Lig1* | 2,16 | 3,849E-02 |
| *Cdk5rap2* | 2,16 | 4,183E-02 |
| *Cd55* | 2,15 | 3,203E-02 |
| *Ehd3* | 2,15 | 1,916E-02 |
| *Oip5* | 2,15 | 3,958E-02 |
| *Tmem51* | 2,14 | 2,121E-02 |
| *Csf1* | 2,14 | 2,554E-02 |
| *Nrp2* | 2,13 | 2,203E-02 |
| *Add3* | 2,12 | 5,894E-04 |
| *Msrb3* | 2,10 | 3,627E-03 |
| *Stk17b* | 2,10 | 7,051E-03 |
| *Wfdc2* | 2,10 | 3,461E-02 |
| *Ahnak* | 2,09 | 1,099E-02 |
| *Nipa1* | 2,05 | 3,570E-02 |
| *Erbb4* | 2,03 | 1,269E-02 |
| *Lgmn* | 2,01 | 2,211E-05 |
| *Smc2* | 2,01 | 2,821E-02 |
| *Il10rb* | 2,01 | 4,717E-03 |
| *Arhgap18* | 1,99 | 1,217E-03 |
| *Gstm3* | 1,98 | 3,633E-02 |
| *Pdzrn3* | 1,98 | 4,221E-02 |
| *Pawr* | 1,98 | 1,507E-02 |
| *Tm4sf4* | 1,97 | 1,055E-03 |
| *Maf* | 1,94 | 5,289E-03 |
| *Txndc16* | 1,93 | 3,684E-02 |
| *Il17rb* | 1,91 | 1,804E-02 |
| *Tmem164* | 1,89 | 7,854E-03 |
| *Sorbs3* | 1,89 | 1,080E-03 |
| *Prim1* | 1,88 | 3,543E-02 |
| *Lamc1* | 1,85 | 1,195E-02 |
| *Slc6a8* | 1,85 | 4,834E-02 |
| *Rnf145* | 1,85 | 3,246E-02 |
| *Tle4* | 1,84 | 8,687E-03 |
| *Ehd4* | 1,84 | 2,574E-02 |
| *Snapc3* | 1,82 | 4,183E-02 |
| *Srgap2* | 1,82 | 2,732E-02 |
| *Nabp1* | 1,82 | 9,578E-03 |
| *Otud7b* | 1,81 | 1,392E-02 |
| *Ano10* | 1,81 | 1,507E-02 |
| *Micu2* | 1,80 | 1,100E-04 |
| *Mgst3* | 1,80 | 1,661E-02 |
| *Scamp5* | 1,79 | 2,166E-02 |
| *Faap20* | 1,77 | 9,273E-03 |
| *E330009J07Rik* | 1,76 | 4,542E-02 |
| *Lmo7* | 1,75 | 3,445E-02 |
| *Mast3* | 1,75 | 1,770E-02 |
| *Hexa* | 1,74 | 1,002E-02 |
| *Armcx3* | 1,74 | 1,455E-02 |
| *Tm6sf2* | 1,73 | 7,489E-03 |
| *Mknk2* | 1,69 | 3,958E-02 |
| *Trak2* | 1,69 | 2,823E-02 |
| *Pik3ap1* | 1,67 | 3,349E-02 |
| *Gnai2* | 1,62 | 4,628E-02 |
| *Hpx* | 1,60 | 2,720E-03 |
| *Tmem176b* | 1,60 | 2,427E-02 |
| *Ctps2* | 1,60 | 4,040E-02 |
| *Plpp2* | 1,58 | 2,554E-02 |
| *Dhfr* | 1,56 | 3,627E-03 |
| *Sult1a1* | 1,53 | 3,879E-02 |
| *Ahsg* | 1,50 | 2,927E-02 |
| *Kng2* | 1,50 | 3,579E-02 |
| *Pdia5* | 1,42 | 4,542E-02 |
| *Acox2* | 0,69 | 2,893E-02 |
| *Lrrc58* | 0,69 | 3,612E-02 |
| *Vwa8* | 0,69 | 3,788E-02 |
| *Dpy19l1* | 0,67 | 4,889E-02 |
| *Ikbkg* | 0,67 | 4,620E-02 |
| *Depdc7* | 0,67 | 4,094E-02 |
| *Ces3b* | 0,66 | 5,880E-03 |
| *Ehhadh* | 0,66 | 1,869E-02 |
| *Hdhd3* | 0,66 | 3,234E-02 |
| *Usf3* | 0,66 | 2,893E-02 |
| *Paqr9* | 0,65 | 3,311E-02 |
| *Gphn* | 0,65 | 1,713E-02 |
| *Tars* | 0,65 | 4,868E-02 |
| *Acox1* | 0,65 | 3,872E-02 |
| *Irf2bp2* | 0,65 | 1,901E-02 |
| *Lactb2* | 0,65 | 2,823E-02 |
| *Abcd3* | 0,65 | 1,247E-02 |
| *Acsl1* | 0,64 | 1,713E-02 |
| *Mbnl1* | 0,64 | 2,848E-02 |
| *Mreg* | 0,63 | 4,820E-02 |
| *Ephx2* | 0,63 | 1,764E-03 |
| *Tcp11l2* | 0,62 | 2,559E-02 |
| *Smim10l1* | 0,62 | 3,311E-02 |
| *Slc22a30* | 0,62 | 4,810E-02 |
| *Ddhd1* | 0,61 | 1,614E-02 |
| *Mpv17l* | 0,61 | 3,248E-02 |
| *Plekhb1* | 0,61 | 2,099E-02 |
| *Pctp* | 0,61 | 3,534E-03 |
| *Akr1c14* | 0,60 | 1,070E-02 |
| *Dnajb2* | 0,59 | 1,559E-02 |
| *Mlxipl* | 0,59 | 3,128E-02 |
| *Tec* | 0,58 | 2,255E-02 |
| *Metap1d* | 0,58 | 4,183E-02 |
| *Ahcyl2* | 0,58 | 4,391E-02 |
| *Crat* | 0,58 | 1,810E-02 |
| *Cyp8b1* | 0,58 | 2,706E-02 |
| *Tcaim* | 0,58 | 8,431E-03 |
| *Chuk* | 0,57 | 2,358E-02 |
| *Chpf2* | 0,57 | 4,763E-02 |
| *Mfsd4b1* | 0,57 | 4,440E-02 |
| *Ido2* | 0,57 | 4,428E-04 |
| *Mal2* | 0,56 | 9,865E-03 |
| *Rnf125* | 0,56 | 1,055E-02 |
| *Gm17753* | 0,56 | 1,292E-02 |
| *Slc17a3* | 0,56 | 7,704E-04 |
| *Chpt1* | 0,56 | 2,632E-03 |
| *Dym* | 0,56 | 4,972E-02 |
| *2310022B05Rik* | 0,55 | 1,580E-02 |
| *Celsr1* | 0,55 | 3,756E-02 |
| *Cyp2d40* | 0,55 | 1,150E-02 |
| *Reep6* | 0,55 | 1,859E-03 |
| *Mup3* | 0,55 | 8,797E-04 |
| *Pnkd* | 0,55 | 2,768E-03 |
| *Dusp1* | 0,55 | 2,674E-02 |
| *Me1* | 0,55 | 1,217E-03 |
| *Slc22a7* | 0,55 | 4,954E-02 |
| *Sult2a8* | 0,54 | 2,441E-02 |
| *Car3* | 0,54 | 9,410E-03 |
| *Tmem19* | 0,54 | 5,542E-06 |
| *Fam96b* | 0,54 | 6,270E-03 |
| *Abcg2* | 0,54 | 9,660E-05 |
| *Retsat* | 0,53 | 1,782E-03 |
| *Ddc* | 0,53 | 5,066E-03 |
| *0610031O16Rik* | 0,53 | 2,512E-02 |
| *Ces1e* | 0,52 | 1,368E-03 |
| *Zfand4* | 0,52 | 1,706E-03 |
| *Fam126b* | 0,52 | 1,109E-02 |
| *Lrfn3* | 0,52 | 1,769E-02 |
| *Gm35986* | 0,52 | 3,958E-02 |
| *Dio1* | 0,51 | 1,328E-04 |
| *Leap2* | 0,51 | 2,539E-02 |
| *Efna1* | 0,51 | 3,609E-02 |
| *Ces2c* | 0,51 | 4,822E-02 |
| *Ttc39c* | 0,51 | 1,360E-02 |
| *Cyp2d9* | 0,50 | 1,070E-02 |
| *Slco1a1* | 0,50 | 1,859E-03 |
| *Hspa13* | 0,49 | 3,908E-02 |
| *Ppdpf* | 0,49 | 4,510E-02 |
| *Cry1* | 0,48 | 1,397E-02 |
| *Spred2* | 0,48 | 2,104E-02 |
| *Snord17* | 0,47 | 2,554E-02 |
| *Sucnr1* | 0,47 | 5,169E-03 |
| *Nr0b2* | 0,47 | 6,307E-03 |
| *Gpd1* | 0,46 | 2,017E-05 |
| *Cyp4a12a* | 0,46 | 5,091E-04 |
| *Thrsp* | 0,45 | 3,591E-03 |
| *Slc17a8* | 0,44 | 6,025E-03 |
| *Mrgprb1* | 0,44 | 3,599E-02 |
| *Fpgs* | 0,44 | 1,123E-02 |
| *Tln2* | 0,43 | 3,485E-02 |
| *Aifm3* | 0,41 | 1,013E-02 |
| *Ppard* | 0,40 | 4,865E-04 |
| *Npas2* | 0,40 | 1,478E-02 |
| *Sult5a1* | 0,40 | 3,636E-02 |
| *Snx29* | 0,40 | 3,627E-03 |
| *Cela1* | 0,38 | 6,543E-06 |
| *Srgap3* | 0,37 | 1,680E-02 |
| *Cyp46a1* | 0,37 | 1,733E-02 |
| *Mkx* | 0,36 | 1,279E-02 |
| *Mup4* | 0,35 | 1,238E-05 |
| *Serpina1e* | 0,35 | 8,397E-05 |
| *Mup6* | 0,34 | 1,916E-07 |
| *Gm19522* | 0,33 | 5,633E-05 |
| *Slc15a5* | 0,33 | 1,402E-06 |
| *Gm40787* | 0,33 | 6,390E-04 |
| *Ces2b* | 0,32 | 7,949E-04 |
| *Mup21* | 0,32 | 4,684E-09 |
| *Rfx4* | 0,32 | 1,113E-05 |
| *Rarres1* | 0,32 | 1,625E-09 |
| *Susd4* | 0,31 | 1,489E-06 |
| *Gm4956* | 0,31 | 3,048E-03 |
| *Gm13152* | 0,31 | 1,131E-04 |
| *Ces4a* | 0,31 | 9,386E-07 |
| *Gm12718* | 0,30 | 1,964E-03 |
| *Extl1* | 0,30 | 5,089E-04 |
| *Sdr9c7* | 0,30 | 1,486E-15 |
| *Serpine2* | 0,27 | 9,292E-05 |
| *Slc13a2* | 0,26 | 2,448E-04 |
| *Capn8* | 0,25 | 4,429E-05 |
| *Dpy19l3* | 0,24 | 2,384E-06 |
| *Hsd3b5* | 0,23 | 2,297E-18 |
| *Cyp7b1* | 0,17 | 6,567E-14 |
| *Elovl3* | 0,16 | 9,860E-16 |
| *Pitx3* | 0,14 | 6,206E-12 |

**Table S2.** Genes differentially expressed in Mcpip1^fl/fl^ and Mcpip1^AlbKO^ hepatocytes collected from 24 weeks old mice.

| **Gene** | **Fold change** | **P value** |
| --- | --- | --- |
| *Lpin1* | 2,24 | 0,0083 |
| *Tmc7* | 1,83 | 0,0397 |
| *Lrp6* | 1,55 | 0,0343 |
| *Lect2* | 0,54 | 0,0343 |
| *Cela1* | 0,52 | 0,0010 |
| *Ppdpf* | 0,52 | 0,0083 |
| *Pla2g16* | 0,49 | 0,0343 |
| *Asap1* | 0,47 | 0,0333 |

**Table S3.** Genes differentially expressed in Mcpip1^AlbKO^ hepatocytes collected from 6 and 24 weeks old mice.

| **Gene** | **Fold change** | **P value** |
| --- | --- | --- |
| *D17H6S56E-5* | 40,69 | 5,641E-49 |
| *Pls1* | 25,73 | 3,520E-29 |
| *Ly6d* | 20,72 | 2,291E-35 |
| *Cenpf* | 20,14 | 1,999E-26 |
| *Mki67* | 19,93 | 3,230E-22 |
| *Cdc20* | 19,46 | 1,212E-27 |
| *Prc1* | 17,82 | 9,646E-21 |
| *Mmp9* | 16,05 | 6,367E-21 |
| *Birc5* | 15,61 | 2,783E-26 |
| *Nuf2* | 15,56 | 3,987E-18 |
| *Ccna2* | 15,00 | 5,060E-26 |
| *Ube2c* | 13,98 | 7,388E-22 |
| *Slc22a26* | 12,29 | 3,407E-14 |
| *Kif20a* | 11,81 | 2,634E-35 |
| *Hmmr* | 11,56 | 9,621E-16 |
| *Racgap1* | 11,27 | 3,695E-14 |
| *Cdkn3* | 11,08 | 3,398E-14 |
| *Cyp2c55* | 10,83 | 9,598E-23 |
| *Cyp2a22* | 10,64 | 5,514E-14 |
| *Hist1h1b* | 10,61 | 1,150E-14 |
| *Top2a* | 10,09 | 7,297E-13 |
| *Igf2bp3* | 10,05 | 9,791E-13 |
| *Tpx2* | 10,05 | 5,483E-12 |
| *Knstrn* | 9,67 | 7,482E-14 |
| *Cib3* | 9,65 | 7,417E-30 |
| *Kif2c* | 9,55 | 1,215E-11 |
| *Ckap2* | 9,51 | 2,811E-11 |
| *Ncapg* | 9,47 | 3,645E-12 |
| *Kifc1* | 9,29 | 2,972E-11 |
| *Hist1h2ak* | 9,20 | 4,000E-12 |
| *Cyp3a44* | 9,09 | 2,884E-11 |
| *Cdk1* | 8,98 | 3,792E-15 |
| *Nusap1* | 8,98 | 1,941E-11 |
| *Aurka* | 8,98 | 1,253E-16 |
| *Cenpe* | 8,84 | 1,820E-10 |
| *Nt5e* | 8,70 | 2,987E-12 |
| *Shcbp1* | 8,38 | 3,371E-12 |
| *Cyp3a11* | 8,30 | 1,789E-10 |
| *Ckap2l* | 8,15 | 1,941E-11 |
| *Tuba8* | 8,15 | 2,960E-25 |
| *Oip5* | 7,82 | 4,984E-12 |
| *Casc5* | 7,81 | 3,341E-09 |
| *Scd2* | 7,71 | 1,884E-13 |
| *Iqgap3* | 7,67 | 1,371E-09 |
| *Cyp2b10* | 7,63 | 4,662E-09 |
| *Cxcr4* | 7,56 | 1,203E-09 |
| *Kif20b* | 7,37 | 3,463E-11 |
| *Cd63* | 7,25 | 3,319E-20 |
| *Ccnb2* | 7,18 | 8,160E-12 |
| *Slfn4* | 7,10 | 3,012E-08 |
| *Arhgap11a* | 7,09 | 9,861E-11 |
| *Bicc1* | 7,06 | 3,362E-09 |
| *Raet1d* | 7,00 | 5,115E-11 |
| *Plk1* | 6,93 | 2,854E-09 |
| *Neurl1b* | 6,86 | 1,506E-08 |
| *Mfge8* | 6,84 | 2,580E-12 |
| *Prom1* | 6,77 | 6,558E-08 |
| *Kif11* | 6,72 | 1,630E-08 |
| *Pbk* | 6,69 | 8,006E-08 |
| *Gm11832* | 6,55 | 3,822E-09 |
| *Cdca2* | 6,44 | 1,311E-07 |
| *Sybu* | 6,31 | 2,438E-07 |
| *Cyp3a59* | 6,31 | 2,326E-12 |
| *Aurkb* | 6,20 | 1,110E-07 |
| *Clec4d* | 6,10 | 5,271E-07 |
| *Cd36* | 6,06 | 7,444E-12 |
| *Lpl* | 6,00 | 2,474E-10 |
| *Trim59* | 5,87 | 9,288E-07 |
| *Cdca5* | 5,83 | 7,524E-08 |
| *Dlgap5* | 5,83 | 1,014E-06 |
| *Sgol2a* | 5,80 | 7,404E-07 |
| *Gpx7* | 5,79 | 7,517E-08 |
| *Rbp1* | 5,69 | 3,060E-15 |
| *Krt23* | 5,61 | 9,436E-08 |
| *Slpi* | 5,60 | 9,936E-07 |
| *Kif22* | 5,59 | 1,966E-06 |
| *Anxa2* | 5,58 | 1,327E-07 |
| *Cdca3* | 5,56 | 1,255E-09 |
| *Epdr1* | 5,39 | 4,420E-06 |
| *Cep55* | 5,32 | 3,222E-06 |
| *Spdl1* | 5,25 | 1,097E-06 |
| *Slc39a4* | 5,24 | 6,755E-18 |
| *Hist1h2bb* | 5,20 | 7,203E-08 |
| *Kif14* | 5,18 | 8,436E-06 |
| *9030619P08Rik* | 5,13 | 1,450E-08 |
| *Uhrf1* | 5,11 | 4,474E-06 |
| *Lrtm2* | 5,09 | 1,127E-05 |
| *Csrp1* | 5,08 | 1,207E-09 |
| *Pla2g16* | 5,00 | 3,673E-13 |
| *Bub1b* | 5,00 | 5,534E-06 |
| *Aspm* | 4,99 | 1,086E-05 |
| *Zc2hc1a* | 4,97 | 1,481E-05 |
| *Arhgap19* | 4,95 | 2,968E-06 |
| *Ttc39a* | 4,88 | 3,212E-06 |
| *Hist1h2ab* | 4,86 | 1,631E-08 |
| *Cxcr2* | 4,79 | 2,995E-05 |
| *Parpbp* | 4,72 | 3,612E-05 |
| *Gipc2* | 4,72 | 6,368E-06 |
| *Sdk1* | 4,71 | 2,570E-05 |
| *S100a8* | 4,68 | 4,296E-05 |
| *Nipal1* | 4,67 | 3,223E-05 |
| *Rad51* | 4,67 | 1,282E-05 |
| *Mcm5* | 4,63 | 3,198E-06 |
| *Fanca* | 4,62 | 3,552E-05 |
| *Kif4* | 4,59 | 4,397E-05 |
| *Cdca8* | 4,59 | 7,872E-06 |
| *Tceal8* | 4,56 | 9,770E-13 |
| *LOC108167426* | 4,52 | 6,869E-05 |
| *Tacc3* | 4,47 | 1,909E-06 |
| *Fignl1* | 4,47 | 8,065E-05 |
| *Sell* | 4,43 | 6,318E-05 |
| *Cyp2c69* | 4,43 | 7,311E-05 |
| *S100a9* | 4,42 | 9,442E-05 |
| *Kif23* | 4,40 | 4,941E-05 |
| *Ahnak* | 4,39 | 3,710E-10 |
| *Car2* | 4,39 | 6,727E-09 |
| *Acss3* | 4,37 | 5,895E-07 |
| *Fgd1* | 4,37 | 2,355E-05 |
| *Kif18b* | 4,36 | 6,332E-05 |
| *Armcx4* | 4,34 | 4,169E-05 |
| *Mis18bp1* | 4,28 | 1,393E-04 |
| *Cyp2a4* | 4,26 | 1,282E-04 |
| *Prtn3* | 4,24 | 1,521E-04 |
| *Gabrb3* | 4,24 | 1,086E-05 |
| *Bhlhb9* | 4,24 | 2,732E-05 |
| *Slco1a4* | 4,23 | 1,686E-04 |
| *Mcm6* | 4,23 | 3,514E-06 |
| *Ttk* | 4,23 | 1,536E-04 |
| *Il1rn* | 4,18 | 4,555E-05 |
| *Arhgef39* | 4,18 | 1,993E-04 |
| *Fam83d* | 4,12 | 2,355E-05 |
| *Nid1* | 4,11 | 4,797E-06 |
| *Gstm3* | 4,10 | 3,709E-10 |
| *Slfn1* | 4,09 | 2,450E-04 |
| *Fam19a2* | 4,09 | 2,555E-04 |
| *1700112E06Rik* | 4,05 | 1,749E-04 |
| *Cand2* | 4,05 | 1,597E-04 |
| *Zwilch* | 4,04 | 2,349E-04 |
| *Anln* | 3,99 | 2,225E-04 |
| *Ube2t* | 3,99 | 6,195E-05 |
| *Asf1b* | 3,97 | 1,686E-04 |
| *Hebp2* | 3,94 | 2,225E-04 |
| *Ucp2* | 3,94 | 4,644E-08 |
| *Mmd2* | 3,92 | 1,189E-05 |
| *Dst* | 3,91 | 3,971E-06 |
| *Tmeff2* | 3,90 | 9,815E-05 |
| *Hpgd* | 3,90 | 2,312E-07 |
| *Cybb* | 3,89 | 2,774E-04 |
| *Fads3* | 3,87 | 1,825E-04 |
| *Hapln1* | 3,85 | 6,121E-04 |
| *Abcd2* | 3,85 | 1,947E-07 |
| *Neil3* | 3,84 | 5,439E-04 |
| *Slfn2* | 3,84 | 3,822E-04 |
| *Slc16a5* | 3,79 | 1,773E-04 |
| *Erbb4* | 3,78 | 7,954E-07 |
| *Mastl* | 3,78 | 2,408E-04 |
| *Ect2* | 3,75 | 6,281E-05 |
| *Glipr2* | 3,74 | 7,392E-04 |
| *Clec2h* | 3,72 | 1,239E-12 |
| *Psrc1* | 3,71 | 9,944E-04 |
| *Ccdc120* | 3,66 | 2,408E-04 |
| *Melk* | 3,61 | 1,374E-03 |
| *Atp6v0d2* | 3,60 | 1,668E-04 |
| *Incenp* | 3,59 | 1,325E-04 |
| *Mad2l1* | 3,56 | 2,877E-06 |
| *2210013O21Rik* | 3,55 | 3,781E-04 |
| *Abcc3* | 3,53 | 1,796E-08 |
| *Ankle1* | 3,53 | 1,479E-03 |
| *Tnfrsf11b* | 3,53 | 1,814E-03 |
| *Tcf24* | 3,52 | 1,256E-03 |
| *Smc2* | 3,51 | 3,634E-06 |
| *Itga6* | 3,47 | 6,216E-05 |
| *Lcn2* | 3,46 | 9,304E-06 |
| *Ugt2b37* | 3,44 | 3,442E-04 |
| *Fanci* | 3,44 | 2,425E-03 |
| *Mfsd7c* | 3,43 | 2,108E-05 |
| *Smpd3* | 3,41 | 1,504E-03 |
| *Sirpa* | 3,39 | 1,043E-03 |
| *Sparc* | 3,38 | 1,972E-04 |
| *H2-Q1* | 3,38 | 7,188E-05 |
| *Stil* | 3,38 | 3,033E-03 |
| *Prr11* | 3,37 | 2,922E-03 |
| *Gcnt1* | 3,36 | 3,104E-03 |
| *Lsp1* | 3,34 | 1,970E-03 |
| *Mcm3* | 3,33 | 8,713E-05 |
| *Pygb* | 3,32 | 9,744E-04 |
| *Elf3* | 3,31 | 1,794E-03 |
| *Hist1h3d* | 3,29 | 1,527E-03 |
| *Abhd2* | 3,28 | 7,014E-16 |
| *Coro1a* | 3,28 | 2,660E-03 |
| *Lysmd2* | 3,28 | 3,968E-03 |
| *Gtse1* | 3,27 | 1,775E-03 |
| *Agpat9* | 3,26 | 1,234E-04 |
| *Hmgcs1* | 3,25 | 5,338E-05 |
| *Snhg18* | 3,24 | 1,536E-04 |
| *Emb* | 3,22 | 4,482E-03 |
| *Vim* | 3,22 | 8,591E-05 |
| *Cd276* | 3,21 | 1,391E-07 |
| *Mcemp1* | 3,21 | 4,475E-03 |
| *Ccdc34* | 3,21 | 1,681E-04 |
| *Alox5ap* | 3,20 | 5,057E-03 |
| *Rad51b* | 3,19 | 4,342E-03 |
| *Col4a2* | 3,18 | 4,936E-04 |
| *Sgol1* | 3,18 | 5,267E-03 |
| *Spsb4* | 3,17 | 5,911E-03 |
| *Uap1l1* | 3,17 | 7,326E-05 |
| *Ncf1* | 3,17 | 3,745E-03 |
| *Stk17b* | 3,14 | 1,909E-06 |
| *Retnlg* | 3,14 | 5,057E-03 |
| *Hexb* | 3,14 | 2,118E-08 |
| *Trem1* | 3,13 | 5,309E-03 |
| *Srgn* | 3,13 | 1,912E-03 |
| *Ercc6l* | 3,13 | 6,833E-03 |
| *Plk4* | 3,12 | 6,959E-04 |
| *Anxa1* | 3,12 | 6,508E-03 |
| *Il16* | 3,09 | 6,886E-03 |
| *Nlrp3* | 3,09 | 7,952E-03 |
| *Gm30784* | 3,08 | 1,668E-04 |
| *Nek2* | 3,07 | 1,014E-05 |
| *Ska1* | 3,06 | 6,680E-03 |
| *Tmem139* | 3,06 | 8,925E-03 |
| *Nsl1* | 3,05 | 6,789E-03 |
| *Mvb12b* | 3,05 | 2,982E-03 |
| *Pglyrp1* | 3,05 | 8,313E-03 |
| *Sirpb1b* | 3,05 | 8,009E-03 |
| *Tceal1* | 3,05 | 5,387E-03 |
| *Pak6* | 3,04 | 7,917E-03 |
| *Rac2* | 3,04 | 9,342E-03 |
| *Nfam1* | 3,04 | 7,644E-03 |
| *Shc2* | 3,04 | 6,886E-03 |
| *Pdp1* | 3,03 | 4,759E-03 |
| *Rad54l* | 3,03 | 9,278E-03 |
| *Tubb6* | 3,03 | 1,239E-06 |
| *Dcaf12l1* | 3,02 | 7,023E-05 |
| *Cenpk* | 3,01 | 5,701E-03 |
| *Ncapg2* | 3,01 | 1,024E-02 |
| *Fgl2* | 3,00 | 1,039E-02 |
| *Ednrb* | 3,00 | 5,057E-03 |
| *Rtn4* | 2,99 | 8,360E-11 |
| *Cd101* | 2,99 | 7,896E-03 |
| *Ska3* | 2,99 | 9,769E-03 |
| *Ly6a* | 2,99 | 5,494E-04 |
| *Cyp2c54* | 2,98 | 5,042E-04 |
| *Gbp9* | 2,97 | 5,518E-03 |
| *Rragd* | 2,97 | 6,434E-03 |
| *Mest* | 2,93 | 1,323E-02 |
| *Panx1* | 2,92 | 1,323E-02 |
| *Gm13056* | 2,92 | 1,373E-02 |
| *Pde4d* | 2,92 | 1,267E-03 |
| *Msantd3* | 2,91 | 1,399E-02 |
| *Hells* | 2,91 | 9,424E-03 |
| *Tagln2* | 2,90 | 5,437E-03 |
| *Pdgfrb* | 2,89 | 1,331E-02 |
| *Elovl7* | 2,89 | 1,087E-02 |
| *Hist1h2ac* | 2,89 | 1,624E-03 |
| *Cd52* | 2,88 | 9,084E-03 |
| *Nckap1l* | 2,88 | 1,206E-02 |
| *Serpinh1* | 2,88 | 8,836E-03 |
| *Il1f9* | 2,88 | 1,546E-02 |
| *Spp1* | 2,87 | 1,557E-02 |
| *Nrm* | 2,87 | 7,482E-03 |
| *Nrep* | 2,87 | 6,142E-04 |
| *Ptprc* | 2,86 | 1,425E-02 |
| *Hist1h1a* | 2,86 | 1,176E-03 |
| *Selplg* | 2,86 | 1,512E-02 |
| *Gm35164* | 2,85 | 1,243E-02 |
| *Npl* | 2,85 | 1,666E-02 |
| *Arhgdib* | 2,85 | 1,101E-03 |
| *Prr15l* | 2,85 | 1,399E-02 |
| *Fmnl1* | 2,85 | 1,689E-02 |
| *Pparg* | 2,84 | 5,154E-03 |
| *Kntc1* | 2,84 | 1,751E-02 |
| *Prlr* | 2,84 | 2,645E-09 |
| *Gja10* | 2,83 | 1,806E-02 |
| *Ncald* | 2,82 | 1,673E-05 |
| *Arhgap30* | 2,82 | 1,666E-02 |
| *Slc25a4* | 2,81 | 6,100E-04 |
| *Cep72* | 2,81 | 1,897E-02 |
| *Ngfrap1* | 2,80 | 1,915E-02 |
| *Cd14* | 2,80 | 1,344E-02 |
| *Cyp26a1* | 2,80 | 4,063E-03 |
| *Pvt1* | 2,80 | 6,790E-03 |
| *Elmo1* | 2,79 | 1,037E-02 |
| *Ccl2* | 2,79 | 2,046E-02 |
| *Nipa1* | 2,79 | 4,539E-04 |
| *BC030867* | 2,78 | 1,548E-02 |
| *Src* | 2,76 | 2,203E-02 |
| *Col1a2* | 2,75 | 2,309E-02 |
| *Slc9a9* | 2,75 | 1,119E-02 |
| *Pdzrn3* | 2,75 | 2,089E-02 |
| *Tinag* | 2,73 | 2,479E-02 |
| *Npdc1* | 2,73 | 8,910E-04 |
| *Npnt* | 2,73 | 1,772E-02 |
| *Rfc4* | 2,72 | 4,865E-04 |
| *Csf3r* | 2,71 | 2,689E-02 |
| *Cd300lf* | 2,71 | 2,727E-02 |
| *Mtmr11* | 2,71 | 2,653E-02 |
| *Diaph3* | 2,71 | 2,724E-02 |
| *Serpinb6a* | 2,70 | 8,587E-05 |
| *Col1a1* | 2,70 | 2,651E-02 |
| *Veph1* | 2,70 | 2,676E-02 |
| *Phlda3* | 2,70 | 2,783E-02 |
| *Prickle2* | 2,69 | 2,871E-02 |
| *Hist4h4* | 2,68 | 8,020E-03 |
| *Fmo2* | 2,68 | 1,719E-02 |
| *Mcm4* | 2,68 | 7,878E-03 |
| *Ntf3* | 2,67 | 2,994E-02 |
| *Mybl1* | 2,67 | 3,052E-02 |
| *4930452B06Rik* | 2,67 | 2,241E-02 |
| *Cyp2c39* | 2,66 | 5,057E-03 |
| *Arhgap44* | 2,66 | 1,074E-02 |
| *Myb* | 2,66 | 2,792E-02 |
| *Msmo1* | 2,65 | 3,597E-04 |
| *Apbb1ip* | 2,65 | 3,064E-02 |
| *Terc* | 2,64 | 3,388E-03 |
| *Bmper* | 2,64 | 3,270E-02 |
| *Slc9a7* | 2,64 | 2,698E-02 |
| *Smc4* | 2,64 | 5,534E-06 |
| *Cyp4a10* | 2,64 | 2,422E-02 |
| *Tlr12* | 2,63 | 9,293E-03 |
| *Syngr4* | 2,62 | 3,226E-02 |
| *Jazf1* | 2,62 | 2,429E-02 |
| *Ces1g* | 2,62 | 2,883E-08 |
| *Mdm1* | 2,62 | 2,994E-02 |
| *Ncapd2* | 2,61 | 3,654E-03 |
| *Vsig10* | 2,61 | 3,105E-02 |
| *2810417H13Rik* | 2,61 | 3,693E-02 |
| *Samsn1* | 2,60 | 3,428E-02 |
| *Entpd1* | 2,60 | 3,000E-02 |
| *Anxa5* | 2,59 | 5,514E-14 |
| *Adgrg1* | 2,59 | 1,323E-02 |
| *Cenpa* | 2,58 | 3,359E-04 |
| *Tnfrsf10b* | 2,58 | 1,915E-02 |
| *Ncf2* | 2,57 | 4,018E-02 |
| *Gm1966* | 2,56 | 3,849E-02 |
| *Fbn1* | 2,55 | 3,794E-02 |
| *Olfr1034* | 2,55 | 3,394E-03 |
| *Rad54b* | 2,55 | 4,213E-02 |
| *Ipcef1* | 2,55 | 2,826E-02 |
| *Ehd4* | 2,54 | 7,199E-05 |
| *Ptp4a3* | 2,53 | 2,773E-02 |
| *Iqgap1* | 2,53 | 8,026E-05 |
| *D130040H23Rik* | 2,53 | 3,129E-02 |
| *Gal3st1* | 2,53 | 2,689E-03 |
| *Gm33543* | 2,53 | 2,596E-05 |
| *Spats2l* | 2,52 | 4,737E-02 |
| *Atp1a3* | 2,52 | 4,424E-02 |
| *Nol4l* | 2,52 | 4,801E-02 |
| *Cenpw* | 2,52 | 3,357E-02 |
| *Traip* | 2,52 | 4,695E-02 |
| *Hist2h4* | 2,51 | 9,586E-03 |
| *Zfp78* | 2,51 | 4,018E-02 |
| *Oasl2* | 2,51 | 1,577E-02 |
| *Cd93* | 2,51 | 1,577E-02 |
| *Gm4952* | 2,50 | 4,472E-04 |
| *Klhl13* | 2,50 | 2,555E-04 |
| *Col4a1* | 2,50 | 1,020E-02 |
| *Elf4* | 2,49 | 4,517E-02 |
| *Fam198b* | 2,49 | 4,672E-02 |
| *Fam25c* | 2,49 | 1,328E-03 |
| *Defb1* | 2,49 | 2,838E-03 |
| *Dsn1* | 2,48 | 4,403E-02 |
| *Zbp1* | 2,48 | 1,918E-02 |
| *Slc22a3* | 2,48 | 1,254E-02 |
| *Amer1* | 2,47 | 4,448E-03 |
| *Itgb2* | 2,47 | 4,671E-02 |
| *Ndrg1* | 2,46 | 3,064E-02 |
| *Alpk1* | 2,46 | 4,001E-02 |
| *Wbp5* | 2,46 | 1,218E-05 |
| *Ces2c* | 2,46 | 4,387E-04 |
| *Dyrk3* | 2,45 | 4,001E-02 |
| *Depdc1b* | 2,43 | 4,018E-02 |
| *Fmo3* | 2,42 | 3,990E-02 |
| *Cyp2c68* | 2,42 | 4,231E-03 |
| *Serpinb8* | 2,42 | 6,458E-03 |
| *Cmpk2* | 2,41 | 6,884E-04 |
| *Cenpq* | 2,39 | 9,767E-03 |
| *Hspg2* | 2,38 | 5,178E-03 |
| *Cenpn* | 2,38 | 3,270E-02 |
| *Slc22a27* | 2,37 | 4,987E-02 |
| *Cenpi* | 2,37 | 4,529E-02 |
| *Abcg1* | 2,37 | 1,963E-02 |
| *Cdkn2c* | 2,37 | 2,369E-04 |
| *Txndc16* | 2,36 | 3,745E-03 |
| *Msn* | 2,35 | 1,689E-02 |
| *Tlr5* | 2,35 | 2,871E-02 |
| *Aldoc* | 2,34 | 5,895E-07 |
| *Pkm* | 2,34 | 3,107E-02 |
| *Ctc1* | 2,32 | 1,789E-04 |
| *Ccl9* | 2,32 | 2,301E-05 |
| *Acot9* | 2,28 | 5,421E-03 |
| *Cyp4f16* | 2,28 | 6,142E-04 |
| *Mis18a* | 2,28 | 2,676E-02 |
| *Syt1* | 2,27 | 9,308E-04 |
| *Vnn3* | 2,27 | 4,606E-05 |
| *Fdft1* | 2,27 | 6,343E-04 |
| *Gstm2* | 2,26 | 4,094E-03 |
| *Kdm6b* | 2,26 | 1,407E-02 |
| *Snrnp25* | 2,25 | 2,976E-02 |
| *Eif4e3* | 2,25 | 4,403E-02 |
| *Dpysl2* | 2,25 | 4,547E-02 |
| *Ubtd2* | 2,25 | 2,758E-02 |
| *Ahr* | 2,24 | 1,775E-03 |
| *Ntrk2* | 2,24 | 3,226E-02 |
| *Hist2h2be* | 2,24 | 1,689E-02 |
| *Aldh1a7* | 2,23 | 2,423E-07 |
| *Spc25* | 2,23 | 3,752E-02 |
| *Hacd1* | 2,22 | 1,997E-03 |
| *Ppm1k* | 2,22 | 9,618E-04 |
| *Bche* | 2,21 | 2,903E-06 |
| *Ptgr1* | 2,21 | 6,508E-03 |
| *Lmnb1* | 2,21 | 4,423E-02 |
| *Mcm2* | 2,21 | 2,698E-02 |
| *Hes1* | 2,19 | 9,293E-03 |
| *Ift57* | 2,19 | 2,676E-02 |
| *Poc1a* | 2,19 | 2,596E-02 |
| *Ank3* | 2,19 | 6,672E-03 |
| *Nt5c3b* | 2,19 | 1,243E-02 |
| *Dab2* | 2,19 | 2,453E-02 |
| *Map3k13* | 2,18 | 2,838E-03 |
| *Rcn2* | 2,17 | 8,109E-06 |
| *Add3* | 2,17 | 1,624E-03 |
| *Slc16a7* | 2,15 | 1,216E-02 |
| *Vnn1* | 2,15 | 2,430E-02 |
| *Tmem237* | 2,15 | 8,290E-03 |
| *Hunk* | 2,13 | 3,805E-02 |
| *Ces2a* | 2,13 | 4,751E-04 |
| *Amot* | 2,13 | 1,237E-02 |
| *Zfp128* | 2,12 | 4,706E-02 |
| *Cep192* | 2,12 | 3,072E-02 |
| *Ifi27l2b* | 2,10 | 1,088E-02 |
| *Apcs* | 2,09 | 2,732E-03 |
| *Dopey2* | 2,08 | 2,422E-02 |
| *Crot* | 2,07 | 6,195E-05 |
| *Scpep1* | 2,07 | 1,668E-04 |
| *Prim1* | 2,06 | 2,313E-02 |
| *Spp2* | 2,06 | 3,609E-03 |
| *Ifi44* | 2,05 | 3,950E-02 |
| *Ermp1* | 2,04 | 4,686E-03 |
| *Slc38a4* | 2,04 | 4,789E-02 |
| *Scarna13* | 2,04 | 1,452E-02 |
| *Antxr2* | 2,03 | 4,378E-03 |
| *Lgals1* | 2,03 | 2,007E-02 |
| *Slc36a4* | 2,03 | 5,832E-03 |
| *Cyp51* | 2,02 | 2,114E-02 |
| *Tm4sf4* | 2,02 | 1,648E-06 |
| *Alg8* | 2,02 | 8,776E-03 |
| *Heg1* | 2,02 | 3,743E-02 |
| *9430020K01Rik* | 2,01 | 4,042E-02 |
| *Dsc2* | 2,00 | 6,315E-03 |
| *Gca* | 2,00 | 4,080E-03 |
| *Pkhd1* | 1,97 | 4,351E-02 |
| *Rpph1* | 1,96 | 1,150E-02 |
| *Gm20594* | 1,95 | 3,270E-02 |
| *Ccdc122* | 1,95 | 2,103E-02 |
| *Vrk2* | 1,94 | 2,309E-02 |
| *Sept11* | 1,93 | 2,114E-02 |
| *Tifa* | 1,92 | 1,494E-03 |
| *Sort1* | 1,92 | 4,482E-03 |
| *Slc1a2* | 1,92 | 4,018E-02 |
| *Slc47a1* | 1,91 | 3,745E-03 |
| *Car5b* | 1,89 | 1,169E-02 |
| *Nsdhl* | 1,89 | 4,536E-02 |
| *Cyp1a2* | 1,88 | 1,055E-02 |
| *Wwox* | 1,88 | 4,365E-02 |
| *Robo1* | 1,87 | 1,024E-02 |
| *Fuca2* | 1,87 | 3,656E-02 |
| *Csrp3* | 1,86 | 7,128E-04 |
| *Ces2e* | 1,86 | 1,328E-03 |
| *Cdh1* | 1,86 | 3,475E-02 |
| *Lcor* | 1,86 | 5,531E-04 |
| *Ctps2* | 1,86 | 1,856E-03 |
| *Fmo1* | 1,82 | 9,424E-03 |
| *Snord15a* | 1,82 | 3,475E-02 |
| *Tmem164* | 1,81 | 1,158E-02 |
| *Lin54* | 1,81 | 3,805E-02 |
| *Inmt* | 1,81 | 4,075E-03 |
| *0610007P14Rik* | 1,80 | 2,373E-02 |
| *Scarna6* | 1,80 | 1,418E-02 |
| *Snord118* | 1,79 | 3,357E-02 |
| *Tm7sf2* | 1,79 | 7,041E-03 |
| *Slc16a1* | 1,79 | 4,678E-02 |
| *E330009J07Rik* | 1,79 | 4,678E-02 |
| *Ifngr1* | 1,78 | 1,642E-02 |
| *Steap2* | 1,77 | 4,403E-02 |
| *Entpd2* | 1,77 | 4,042E-02 |
| *Tmem176b* | 1,76 | 1,585E-03 |
| *Ckap5* | 1,75 | 4,018E-02 |
| *Cetn2* | 1,75 | 2,339E-02 |
| *Ifih1* | 1,73 | 2,922E-03 |
| *Nabp1* | 1,73 | 2,946E-02 |
| *Idh2* | 1,72 | 3,903E-03 |
| *Sdc1* | 1,72 | 1,025E-02 |
| *AW112010* | 1,72 | 7,168E-03 |
| *Cyp3a25* | 1,72 | 3,129E-02 |
| *Ggh* | 1,70 | 4,120E-02 |
| *Nme7* | 1,70 | 3,412E-02 |
| *Micu2* | 1,70 | 2,369E-04 |
| *Gm12942* | 1,69 | 3,475E-02 |
| *Rpa1* | 1,69 | 9,767E-03 |
| *Agmo* | 1,68 | 4,801E-02 |
| *Cxadr* | 1,67 | 1,342E-02 |
| *Tmem218* | 1,67 | 3,569E-02 |
| *Ugt2b1* | 1,66 | 9,278E-03 |
| *Hn1l* | 1,66 | 2,246E-02 |
| *Mcm7* | 1,65 | 1,772E-02 |
| *Ctso* | 1,65 | 6,047E-03 |
| *Arl6ip1* | 1,64 | 3,603E-03 |
| *Slc35e2* | 1,64 | 1,518E-02 |
| *Marveld2* | 1,63 | 2,722E-02 |
| *Ttpa* | 1,63 | 6,120E-03 |
| *Armc8* | 1,63 | 2,871E-02 |
| *Cmbl* | 1,63 | 4,501E-02 |
| *Ahsg* | 1,62 | 1,035E-03 |
| *Pgap2* | 1,62 | 1,775E-03 |
| *Cyp2c44* | 1,62 | 9,631E-03 |
| *Tmtc3* | 1,61 | 4,693E-02 |
| *Steap3* | 1,60 | 4,871E-02 |
| *Dpp7* | 1,60 | 9,077E-03 |
| *Suox* | 1,59 | 1,099E-03 |
| *Vwa5a* | 1,59 | 1,537E-02 |
| *Ilvbl* | 1,59 | 6,789E-03 |
| *Sptssa* | 1,58 | 4,672E-02 |
| *Sdcbp* | 1,56 | 1,835E-02 |
| *Slc4a4* | 1,55 | 6,929E-03 |
| *Rfwd3* | 1,55 | 4,018E-02 |
| *Asah1* | 1,55 | 4,729E-02 |
| *Prune* | 1,54 | 3,226E-02 |
| *Agtr1a* | 1,52 | 1,066E-02 |
| *Dsp* | 1,51 | 3,302E-02 |
| *Hpx* | 1,51 | 1,816E-02 |
| *Phkb* | 1,49 | 4,213E-02 |
| *Cstb* | 1,48 | 3,090E-02 |
| *Calm2* | 1,48 | 1,342E-02 |
| *Dag1* | 1,48 | 2,776E-02 |
| *Ebp* | 1,44 | 4,042E-02 |
| *Ndfip2* | 1,41 | 3,419E-02 |
| *Bsg* | 1,36 | 4,213E-02 |
| *Usf2* | 0,70 | 4,789E-02 |
| *Fgd6* | 0,69 | 4,042E-02 |
| *Hnrnpul2* | 0,68 | 4,536E-02 |
| *Ube2q1* | 0,68 | 3,401E-02 |
| *Dnajc7* | 0,67 | 3,342E-02 |
| *Alkbh5* | 0,67 | 2,526E-02 |
| *Tceb3* | 0,66 | 4,018E-02 |
| *Ahsa2* | 0,66 | 4,776E-02 |
| *St3gal4* | 0,65 | 2,653E-02 |
| *Pdcd4* | 0,65 | 1,498E-02 |
| *Otud5* | 0,64 | 4,858E-02 |
| *Epas1* | 0,64 | 2,822E-02 |
| *Cdip1* | 0,64 | 1,552E-02 |
| *Trim41* | 0,64 | 4,862E-02 |
| *Pnpla6* | 0,63 | 2,994E-02 |
| *Acbd6* | 0,63 | 3,206E-02 |
| *Egln1* | 0,63 | 4,359E-02 |
| *Phf20l1* | 0,62 | 5,309E-03 |
| *Pi4ka* | 0,62 | 3,316E-02 |
| *Mgat2* | 0,62 | 3,707E-02 |
| *Mup3* | 0,61 | 7,168E-03 |
| *Zfp445* | 0,61 | 3,132E-03 |
| *Mavs* | 0,61 | 3,064E-02 |
| *Anks1* | 0,61 | 4,529E-02 |
| *Tmem19* | 0,60 | 1,623E-04 |
| *Sdccag3* | 0,60 | 3,843E-02 |
| *Ifi27* | 0,60 | 3,226E-02 |
| *Ern1* | 0,60 | 3,412E-02 |
| *Ttc7b* | 0,60 | 4,067E-02 |
| *Rnf25* | 0,60 | 3,090E-02 |
| *Elmod3* | 0,60 | 2,972E-02 |
| *Vegfb* | 0,60 | 1,392E-02 |
| *Trib1* | 0,60 | 3,324E-02 |
| *Crebl2* | 0,59 | 3,053E-02 |
| *Brd1* | 0,59 | 1,414E-02 |
| *Hist1h1c* | 0,59 | 4,829E-03 |
| *Scnn1a* | 0,59 | 1,243E-02 |
| *Rufy3* | 0,59 | 3,053E-02 |
| *Secisbp2* | 0,58 | 2,429E-02 |
| *Inpp4a* | 0,58 | 2,733E-02 |
| *Eml3* | 0,58 | 9,767E-03 |
| *Kcnn2* | 0,58 | 5,144E-03 |
| *Cwf19l1* | 0,58 | 2,724E-02 |
| *Fam13b* | 0,58 | 4,424E-02 |
| *Pnisr* | 0,57 | 3,000E-02 |
| *Eef2kmt* | 0,57 | 3,226E-02 |
| *Taok2* | 0,57 | 2,296E-02 |
| *Zc3h7a* | 0,57 | 1,680E-02 |
| *Cd2bp2* | 0,56 | 1,406E-02 |
| *Tkfc* | 0,56 | 1,005E-02 |
| *Gm38426* | 0,56 | 4,147E-02 |
| *Ptov1* | 0,56 | 9,293E-03 |
| *Zfp180* | 0,56 | 1,766E-02 |
| *1110059G10Rik* | 0,56 | 1,719E-02 |
| *Polb* | 0,55 | 2,422E-02 |
| *Zfp598* | 0,55 | 1,546E-02 |
| *Zpr1* | 0,55 | 9,278E-03 |
| *H1f0* | 0,55 | 5,057E-03 |
| *Aars2* | 0,55 | 3,138E-02 |
| *Ppfia1* | 0,54 | 2,053E-03 |
| *Tns1* | 0,54 | 2,309E-02 |
| *Abcg5* | 0,54 | 2,871E-02 |
| *Dhx33* | 0,54 | 3,611E-02 |
| *Cyp2u1* | 0,54 | 2,934E-02 |
| *Ccdc84* | 0,54 | 1,711E-02 |
| *Trabd* | 0,54 | 3,115E-03 |
| *H2-Q6* | 0,53 | 4,612E-02 |
| *Cebpb* | 0,53 | 2,202E-02 |
| *Dvl1* | 0,53 | 3,075E-02 |
| *Spsb2* | 0,53 | 3,047E-02 |
| *Ehbp1l1* | 0,53 | 1,806E-02 |
| *Dusp1* | 0,53 | 6,839E-03 |
| *Mprip* | 0,53 | 1,642E-02 |
| *Srsf4* | 0,53 | 9,586E-03 |
| *Ppp1r3b* | 0,53 | 2,863E-02 |
| *Car5a* | 0,52 | 2,715E-02 |
| *Rabl6* | 0,52 | 4,706E-02 |
| *Junb* | 0,52 | 4,318E-02 |
| *Taf3* | 0,51 | 3,161E-02 |
| *Kansl3* | 0,51 | 4,317E-02 |
| *Tmie* | 0,51 | 2,241E-02 |
| *Khk* | 0,51 | 9,248E-05 |
| *Ppan* | 0,51 | 4,193E-02 |
| *Sdr9c7* | 0,50 | 2,616E-03 |
| *Ppp1r9a* | 0,50 | 2,653E-02 |
| *Nrbp2* | 0,50 | 1,086E-05 |
| *Atg16l2* | 0,50 | 4,998E-02 |
| *Tnk2* | 0,50 | 2,767E-02 |
| *Cela1* | 0,50 | 2,195E-03 |
| *Scd1* | 0,49 | 1,004E-02 |
| *Trak1* | 0,49 | 4,482E-03 |
| *Slc17a8* | 0,49 | 3,822E-02 |
| *Srebf1* | 0,49 | 2,175E-02 |
| *Tmem39a* | 0,49 | 1,246E-02 |
| *Chd3* | 0,49 | 3,357E-02 |
| *Madd* | 0,49 | 4,236E-02 |
| *Camk2b* | 0,48 | 1,546E-02 |
| *Slc9a3r2* | 0,48 | 4,789E-02 |
| *Fndc4* | 0,48 | 7,321E-06 |
| *Plekhb1* | 0,47 | 2,574E-03 |
| *Fgfr3* | 0,47 | 4,051E-02 |
| *Cspg5* | 0,47 | 3,931E-02 |
| *4430402I18Rik* | 0,47 | 4,730E-02 |
| *Tspyl2* | 0,46 | 4,801E-02 |
| *Crygn* | 0,46 | 3,064E-02 |
| *Rfx4* | 0,46 | 4,708E-02 |
| *Gpc1* | 0,46 | 4,178E-02 |
| *Aacs* | 0,45 | 3,273E-02 |
| *Trim39* | 0,45 | 3,475E-02 |
| *Gm39079* | 0,45 | 6,826E-04 |
| *Serpina1e* | 0,45 | 2,733E-02 |
| *Rpp40* | 0,44 | 3,401E-02 |
| *Mapk15* | 0,44 | 3,909E-02 |
| *Abcg8* | 0,44 | 1,828E-03 |
| *Me1* | 0,43 | 4,619E-03 |
| *1700094D03Rik* | 0,43 | 8,289E-04 |
| *Tcof1* | 0,43 | 1,114E-02 |
| *Mug-ps1* | 0,43 | 3,433E-02 |
| *Gtf2ird2* | 0,43 | 4,672E-02 |
| *Slc15a5* | 0,43 | 1,298E-03 |
| *Acot11* | 0,42 | 1,850E-02 |
| *Irf5* | 0,42 | 2,853E-03 |
| *Unc13b* | 0,42 | 4,155E-05 |
| *Gm35986* | 0,41 | 2,425E-03 |
| *Kcnk10* | 0,41 | 4,536E-02 |
| *Gm13152* | 0,40 | 1,281E-02 |
| *Fam20c* | 0,40 | 1,775E-03 |
| *H2-K2* | 0,40 | 5,483E-05 |
| *Cxcl13* | 0,39 | 4,529E-02 |
| *Nox4* | 0,39 | 4,658E-06 |
| *Mlxipl* | 0,39 | 1,218E-05 |
| *Gm31264* | 0,39 | 2,850E-02 |
| *Gna14* | 0,39 | 2,152E-02 |
| *Whrn* | 0,39 | 3,591E-02 |
| *Cyp17a1* | 0,38 | 6,355E-04 |
| *Gck* | 0,38 | 1,328E-03 |
| *Mbd1* | 0,38 | 7,644E-03 |
| *Sarm1* | 0,38 | 3,247E-02 |
| *Pklr* | 0,37 | 3,597E-04 |
| *Celsr1* | 0,37 | 9,816E-09 |
| *Syt3* | 0,37 | 1,750E-02 |
| *Stk19* | 0,36 | 5,982E-07 |
| *Slc2a4* | 0,36 | 2,212E-02 |
| *Mfsd2a* | 0,36 | 3,399E-03 |
| *Dlgap1* | 0,36 | 1,313E-02 |
| *Dusp4* | 0,35 | 1,835E-02 |
| *Nat8* | 0,35 | 1,853E-03 |
| *Lepr* | 0,35 | 3,524E-03 |
| *Pitx3* | 0,33 | 1,024E-02 |
| *Dpy19l3* | 0,33 | 2,192E-03 |
| *Fam193b* | 0,33 | 3,532E-04 |
| *Col5a3* | 0,32 | 3,645E-04 |
| *LOC108169060* | 0,31 | 5,056E-03 |
| *Chpf* | 0,31 | 4,619E-03 |
| *1600002H07Rik* | 0,29 | 5,616E-09 |
| *Serpina4-ps1* | 0,29 | 2,174E-03 |
| *Fgf21* | 0,29 | 9,605E-05 |
| *Sytl1* | 0,29 | 2,407E-06 |
| *Npr2* | 0,29 | 1,437E-14 |
| *Cadm4* | 0,28 | 9,482E-09 |
| *Thrsp* | 0,28 | 8,065E-05 |
| *Acpp* | 0,28 | 3,593E-04 |
| *Tiam2* | 0,28 | 1,410E-06 |
| *Reck* | 0,28 | 7,199E-05 |
| *Bmyc* | 0,27 | 9,110E-05 |
| *Gm4956* | 0,26 | 3,048E-04 |
| *Susd4* | 0,25 | 1,240E-09 |
| *Slc3a1* | 0,25 | 2,913E-20 |
| *Cyp21a1* | 0,24 | 1,782E-04 |
| *Col27a1* | 0,24 | 5,895E-07 |
| *Capn8* | 0,23 | 2,562E-05 |
| *Tmem28* | 0,22 | 7,790E-05 |
| *Hsd3b5* | 0,22 | 2,633E-18 |
| *Chrm3* | 0,21 | 4,511E-06 |
| *Slc13a2* | 0,20 | 3,152E-06 |
| *Gm15343* | 0,20 | 4,037E-11 |
| *Rgs16* | 0,20 | 4,730E-06 |
| *Grm8* | 0,19 | 4,303E-06 |
| *Gm12718* | 0,18 | 2,116E-06 |
| *Cyp7b1* | 0,17 | 5,842E-13 |
| *Srgap3* | 0,16 | 2,512E-08 |
| *Ar* | 0,15 | 1,126E-08 |
| *Mup21* | 0,13 | 8,361E-18 |
| *Extl1* | 0,12 | 1,200E-13 |
| *Trhde* | 0,12 | 9,313E-11 |
| *Serpine2* | 0,04 | 4,685E-42 |
| *Moxd1* | 0,00 | 4,104E-97 |
